# A comparative analysis of fruit feeding among Mediterranean passerine birds

**DOI:** 10.64898/2026.03.20.712853

**Authors:** Pedro Jordano, Elena Quintero, Jorge Isla

## Abstract

Fleshy fruits underpin a major mutualistic pathway linking plants and birds in Mediterranean scrublands, yet we still lack a mechanistic understanding of how ecomorphological and digestive traits constrain fruit use, foraging behaviour, and ultimately the effectiveness of avian seed dispersal. Here we assemble an integrative dataset for 146 Iberian bird species combining external morphology, digestive anatomy, diet composition, and fine-grained observations of fruit foraging and handling obtained from standardized focal watches and camera traps at fruiting plants. We classify species into five functional feeding groups (seed dispersers, pulp consumers, pulp consumer-dispersers, pulp consumer-seed predators, non-frugivores) and ask how suites of traits map onto these feeding modes and onto quantitative metrics of frugivory and feeding rate. Across species, the proportion of diet volume made up by fleshy fruits increases with gape width and faster food transit, and decreases with larger gizzards and longer intestines, indicating a tight coupling between frugivory and traits that enable rapid processing of dilute, pulp-rich food. A small subset of traits (body mass, gape width, gizzard mass, transit time) explains over half of the interspecific variation in fruit consumption, with ecomorphological and digestive characters contributing roughly equally to explained variance. Per-visit feeding rates and numbers of fruits ingested per visit scale positively with body mass, and canonical discriminant analysis reveals distinct multivariate trait syndromes separating seed dispersers from pulp consumers, seed predators, and non-frugivores. These trait syndromes, and the associated differences in handling mode and feeding speed, provide a mechanistic link between individual-level foraging decisions and the sparsity, asymmetry, and effectiveness of plant-frugivore interaction networks in Mediterranean systems. Our results highlight how trait-based constraints shape not only who interacts with whom, but also how efficiently seeds are removed and dispersed across a diverse frugivore assemblage.

## 2 Introduction

Fleshy fruits provide a staple food for many bird species in Mediterranean scrublands, where frugivory is often associated with migratory behavior and timing of migration (Herrera, 1984b, 1987). The fruit food of avian frugivores in the Iberian Peninsula, in particular in Mediterranean habitats, has been profusely documented (e.g., Herrera, 1984a,b; Jordano, 1985, 1987a,b, 1988; Snow and Snow, 1988; Debussche and Isenmann, 1989; Fuentes, 1994, 1995; Guitián et al., 2000; Herrera, 2004; García, 2016; Costa et al., 2020; González-Varo et al., 2021), revealing a key role of these species in the maintenance and persistence of this biome, where their role as seed dispersers represents a fundamental ecological service for forest dynamics (Bascompte and Jordano, 2014; Rebollo et al., 2019). In turn, use of fleshy fruits as food has largely contributed to the diversification and speciation of multiple avian subclades and has a distinct signal in the large-scale biogeography of the whole group (Kissling et al., 2009; Burin et al., 2016; McFadden et al., 2022). The role of frugivorous birds is thus central for the natural regeneration of Mediterranean forests, and a myriad of ecological interactions are embedded in this close mutualistic relationship between the plants and the avian seed dispersers. A central aspect to this type of mutualistic interaction is the effectiveness of fruit foraging by the animals (Schupp et al., 2017), which in turn receive a food resource provisioning from the plants (Quintero et al., 2020). The food rewards represented by the fleshy fruits are frequently obtained by the birds during short feeding bouts while foraging at the fruiting plants, and therefore ample variation exists among frugivore species in their fruit handling behaviors and feeding rates. Understanding such variation is essential to interpret the range of effectiveness in fruit processing that we frequently see in nature when studying avian frugivores.

Previous work focused on local Mediterranean lowland and highland habitats (Herrera, 1984b,a) has documented how the subset of avian frugivore species has distinct features in terms of ecomorphological variation, and digestive traits related to variable fruit consumption. Frugivory is often associated across species with variation in body mass, gape width (closely related to fruit handling abilities), and adaptations of the digestive system involving shorter gut passage time through relatively longer intestines and relatively larger livers that increase the ability to detoxify secondary compounds in the fruit pulp of some species. Such trends of variation are observable even when comparing closely related species in the same family (Jordano, 1987a). Yet, no general overview exists on Mediterranean bird frugivory especially including both non-passerine and passerine frugivores. General reviews for neotropical frugivorous birds (e.g., Moermond and Denslow, 1985; Moermond et al., 1986) and temperate avian frugivores (e.g., Turcek, 1961; Snow and Snow, 1988; Stiebel and Bairlein, 2008) suggest ample variation in fruit foraging and handling modes, as well as in feeding rates, resulting in highly variable reliance in fruit food. Moreover, a central and unexplored aspect of frugivore effectiveness relates to the behavioral correlates of fruit feeding, i.e., whether species strongly relying on fruit food have characteristic foraging behavior features that differ from non-frugivore species.

Frugivorous birds consume an extremely wide range of fleshy-fruited species that differ in colour, size, hardness, accessibility, availability, nutrient content of the pulp, and presence of secondary compounds that act as digestion inhibitors or that simply are toxic (Jordano, 2014). Comparative studies documenting fruit feeding behaviors, however, have reported consistent patterns shared by birds of similar morphologies, suggesting that ecomorphological limitations may severely constrain fruit choice by avian frugivores (e.g., Herrera and Jordano, 1981; Santana and Milligan, 1984; Denslow and Moermond, 1982; Jordano, 1987a; Stiebel and Bairlein, 2008). Such observations argue strongly that a bird’s morphology must provide at least part of the basis for choosing among fruits (Moermond et al., 1986) and underscore the relevance of ecomorphological and foraging data to understand fruit selection by frugivorous animals (González-Varo and Traveset, 2016). Ecomorphological constraints help explain why plant-frugivore interaction networks are so sparse (low density of interactions) (Bascompte and Jordano, 2014), for example when fruit size is so large that the interaction is impossible because size limitations do not allow an efficient handling and consumption of the fruit by the frugivore. These types of size mismatches for instance may explain up to 10% of the missing interactions in plant-frugivore networks (Olesen et al., 2011; Bascompte and Jordano, 2014) and, importantly, may also limit seed dispersal success of the respective plants involved in the interactions (Rehling et al., 2021). Therefore, while digestive adaptations may limit reliance on fruit food, ecomorphological constraints will determine foraging modes, maximum feeding rates, and accessibility conditions when foraging for fruits (Pizo et al., 2021). Ultimately, those are the types of drivers determining fruit preference and variation in frugivory.

Here we overview the diversity of foraging modes and fruit feeding rates of an ample diversity of Mediterranean frugivores, spanning from species relying heavily on fruit food, especially during migration and wintering, to species just using the fruits sporadically or not at all. We first characterize the Mediterranean frugivore avifauna in terms of ecomorphological and physiological traits, then explore variation in fruit use among species and how variable are feeding rates and fruit handling behaviors, and finally discuss the relations between ecomorphological variation across species and variation in reliance on fruit food.

## 3 Material and Methods

Here we report a large dataset of avian frugivore feeding observations and records obtained over the years with a variety of methods. Most of these methods aimed at recording plant-avian frugivore interactions to assess ecological interactions’ biodiversity. These methods are quite diverse and are summarized and reviewed in Quintero et al. (2021); Moracho, E et al. (2026). Here we provide a brief outline of the main methodology used.

### 3.1 Bird morphology

Data on ecomorphological traits were obtained from birds in the hand, usually mist-netted (see details in Jordano, 1985, and references therein) in different localities of Southern Spain (Jordano, 1982; Herrera, 1984b; Jordano, 1985), summarized in Herrera (2004). This totalled 146 species and *N >* 6000 individuals sampled. External morphological traits include: wing length (mm), obtained from folded wing, stretched; tail length (mm); tarsus length (mm); culmen length, to cranium (mm); culmen length, to feathers (mm); culmen width (mm; measured at the distal edge of the nostrils); culmen height (mm; measured at the distal edge of the nostrils); gape width at the mouth commissures (mm, measured with the bill open); body mass (g); gut passage time (min), time to first appearance of barium sulphite marker administered orally to the bird; gizzard mass (g); liver mass (g); intestine length (from pylorus to cloaca)(mm); transit speed of food throughout the digestive system (mm/min), estimated as *intleng/gpt*.

As sources for measurement data we used either measurements of birds in hand (mist-netted) or obtained from specimens deposited at the Estación Biológica de Doñana (CSIC) collection. Some data (see Suppl. Mat.) were obtained from Tobias (2022) and Dunning (2008). Internal anatomical data and gut passage times were obtained as explained in detail in Herrera (1984a); Jordano (1987a). For a complete species list and summary data see Suppl. Mat. Tables 1 and 2. The relative fit between fruit size and gape width of birds is a limiting trait for the range of fruit sizes that can be swallowed and efficiently handled (Levey, 1987). To estimate it we used: 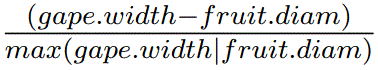, with fruit.diam data obtained from (Valido et al., 2011).

### 3.2 Feeding observations

Observations of birds feeding on fruits at the plants were recorded during timed watches, usually 1 h length, using 10x40 binoculars and/or telescope and two hand chronometers, and spaced at different times of the day (see Herrera and Jordano, 1981; Jordano, 1982; Snow and Snow, 1988; Jordano and Schupp, 2000, for general descriptions of methods for recording feeding rates). We also used camera traps to monitor bird activity at several species (see Quintero et al., 2021; Villalva et al., 2024, for a detailed description). Videos with feeding records were selected and the descriptors of feeding behavior and feeding rates described below were recorded with the same techniques.

Individual plants were selected for focal sampling, either with direct watches or with camera traps, usually following a randomly-stratified protocol to capture the range of natural conditions where fruiting plants are encountered (see Jordano and Schupp, 2000; Villalva et al., 2024, for details). In a given site, individual plants were randomly selected within distinct sectors of the area so as to cover a representative portion of the local conditions, plant size, vegetation cover, edge effects, etc.

We placed variable numbers of camera-traps in each area and plant species (between 10-25) and were rotated periodically among different focal plants. Camera-traps were active for 1–10 days in each focal plant, depending on camera type and plant species (i.e., whether nocturnal mammal frugivores also visited the plants), and then moved to sample other individuals. Cameras were fitted at 1 m above ground and 2-3 m from the focal plant focusing on both ground and the bottom of the plant to detect visits on the plant and from the ground. depending on camera type, the cameras were active day and night or only during the day, in video mode, with 10 seconds per video and 2 seconds between recordings. When using Go-Pro cameras we set to continuous live video recording for 2h. All camera-trap videos containing frugivore visits were recorded. All those detections in which the animal was only walking/flying, or showed a behaviour different from a fruit-foraging visit (e.g., passive perching, hunting prey, scent marking) were not recorded as visitation events.

Visits were recorded since the first sight of the bird when entering the fruiting plant, accumulating the visit duration in the first chronometer (total time of the observation, in s), and following the bird during the whole visit. In some cases, only partial visits we recorded, either because the entering or leaving of the bird could not be recorded, when lost of sight after a feeding sequence at the plant. During the observation of the visit, the second chronometer was used to accumulate just the time the bird was seen stopped, perched with no signs of being actively searching for fruits (time bird was stopped, not moving or feeding, in s). During the feeding bout we recorded the number of fruits “touched”, i.e., with attempts to pick them (Snow and Snow, 1988), the number of fruits swallowed, the number of fruits dropped, and the number of fruits failed to handle and/or ingest, yet not dropped. In addition, we recorded the number of fruits carried away from the tree (e.g., in the bill), the number of fruits pecked (when birds were pecking to tear-off pieces of pulp, either detaching or not the fruit from its pedicel), the number of fruits picked, the number of fruits taken away from tree (swallowed or carried), and the number of fruits estimated to be dropped beneath the plant. We also annotated the number of moves the bird did during the feeding sequence, i.e., location shifts involving jumps or moves between branches or any other displacement. Feeding sequences were summarized from averaged feeding rates across plant species using sawtooth or step graphs (Cody, 1968). These graphs illustrate both the feeding rate and the sequences of active food searching and stops while perched.

We recorded different types of fruit picking behaviors (Suppl. Mat. Fig. 1), including the number of fruits taken while perched from branch, specifying the number of fruits taken below the perched position (while hanging from branch), the number of fruits taken to the front of the perch, or taken upwards from the perched position (Suppl. Mat. Fig. 1). Other feeding manoeuvres recorded included the number of fruits taken while jumping (sallying, picking the fruit on the flight when jumping from a perch to another), while flycatching (bird flying from a perch and stalling in front of the infructescence while picking the fruit), or directly foraging on the ground. Similar categorizations of foraging have been used in similar studies (e.g., Fitzpatrick, 1980; Moermond and Denslow, 1985; Gabriel and Pizo, 2005).

A few main types of feeding types have been described for frugivorous birds (e.g., Levey, 1987; Snow and Snow, 1988), basically separating birds able to swallow fruits whole (SD, swallowers or gulpers), and disperse seeds internally, from those that peck pieces of pulp by tearing-off with repeated pecks at the fruit (PC, pulp consumers, pulp thieves) and drop the seeds (Snow and Snow, 1988; Simmons et al., 2018). Some PC species are able to disperse some seeds, however (PC/SD species Jordano and Schupp, 2000) because they peck the fruit and occasionally ingest seeds and/or carry seeds away from the plant without dropping them. Seed predator species are typically granivorous birds that pick fruits to extract the seeds and crack them open to ingest the seed embryo; in some plant species they may also act as pulp consumers rather than seed predators (we include both in PC/SP) (). Some granivorous species can disperse some seeds, however, especially when the fruit is depulped and the seed incidentally falls to the ground due to handling failures. This behavior may represent up to 30% of seeds dispersed by an avian frugivore assemblage (Heleno et al., 2011) we restrict ourselves to the scenario of PC/SP interactions. Finally, non-frugivorous species (NF) are also considered here, including those that only very occasionally consume some fruit.

The proportion of the diet made up by fleshy fruit was summarized from existing literature (e.g., Guitián, 1983; Herrera, 1984b,a; Jordano, 1984, 1987a,b; Herrera, 2004; Jordano, 1982; Bascompte and Jordano, 2014, and references therein).

### 3.3 Statistical analysis

Data and code for analyses are available (see Suppl. Mat. and Data Availability sections). Analyses on both the ecomorphological and the feeding rates datasets were carried out on standardised, centred variables using the R version 4.2.1 (R Core Team, 2022).

We used a canonical discriminant analysis (CDA) to assess bird ecomorphological traits that maximise the differences between the five major fruit feeding functional groups: seed dispersers (SD), pulp consumers (both PC and pulp consumers/seed dispersers, PC/SD), pulp consumers/seed predators (PC/SP), and non-frugivores (NF). We set ecomorphological traits and feeding traits as predictor variables, and frugivory type (feeding group) affiliation as a categorical dependent variable. CDA allows us to determine if there is a distinct combination of traits that differentiates the feeding groups. In other words, to what extent would we be able to predict the affiliation to a fruit feeding mode (SD, NF, PC, PC/SD, PC/SP) of a bird species based on its ecomorphological and feeding mode traits.

Prior to computing the CDA, we removed correlated variables by conducting a multicollinearity analysis using Variance Inflation Factor (VIF, threshold = 3). Additionally, we identified variable weights on the discriminant function based on the Wilks’ *λ* criterion. For the analysis we used the candisc v. 0.8.6 R package (Friendly and Fox, 2021) and Wilks’ *λ* with the greedy.wilks function in klaR v. 1.7.1 R package (Weihs et al., 2005). We also estimated the 90% equiprobability ellipses to delineate variation trends around the multivariate centroids of each group, as implemented in candisc (Friendly and Fox, 2021). Relationships among fruit feeding and feeding rates and their correlates with ecomorphological traits were tested with linear models, using the R package lme4 (Bates et al., 2015). Additional R packages used include: ade4 v. 1.7.19 (Thioulouse et al., 2018), grateful v. 0.1.11 (Rodríguez-Sánchez et al., 2022), kableExtra v. 1.3.4 (Zhu, 2021), knitr v. 1.39 (Xie, 2022), rmarkdown v. 2.15 (Allaire et al., 2022), and tidyverse v. 1.3.2 (Wickham et al., 2019).

## 4 Results

### 4.1 Patterns of fruit consumption

Avian species potentially interacting with fruits in the Mediterranean range over two orders of magnitude in body mass (Fig. 1; see also Suppl. Mat. Table 1 for the complete list and data). Reliance on fruit food for birds varies accordingly, from species never consuming fruits (e.g., mostly insectivorous muscicapids, most larks and emberizids), those consuming fruit sporadically (e.g., several *Phylloscopus*), frequently (e.g., most *Sylvia* warblers, corvids, and the largest muscicapids), and the strongly frugivorous species (e.g., largest *Sylvia*, the thrushes, and *Bombycilla*). The percentage of the diet volume made up by fleshy fruits (*frupv*, Table 1, Suppl. Mat. Table 1) indicates the overall relative importance of fruits in relation to other food resources in the diet. Yet another component is how extensive is frugivory among individual birds in a given locality or season. Most species with high *frupv* also show extensive consumption of fleshy fruits: the fruits can be found in a large fraction of samples from different individuals. In general, fruit food is supplemented with arthropods and/or seeds in the case of granivorous species (e.g., finches, buntings, partridge) or vegetative plant parts (leaves; pigeons, etc.) or even carrion (corvids). Fig. 1 includes more than strictly Mediterranean species just to encompass the whole diversity of avian frugivores in the Iberian Peninsula. Across species, the percentage of fruit food (% diet volume contributed by fleshy fruits) positively and significantly correlates with body mass (*r* = 0.3209, *F* = 16.40, *P* = 8.2*e^−^*^5^, *d.f.* = 1, 143; frupv= -0.0763 + 0.0728[body mass], on log-transformed data).

**Figure 1:**
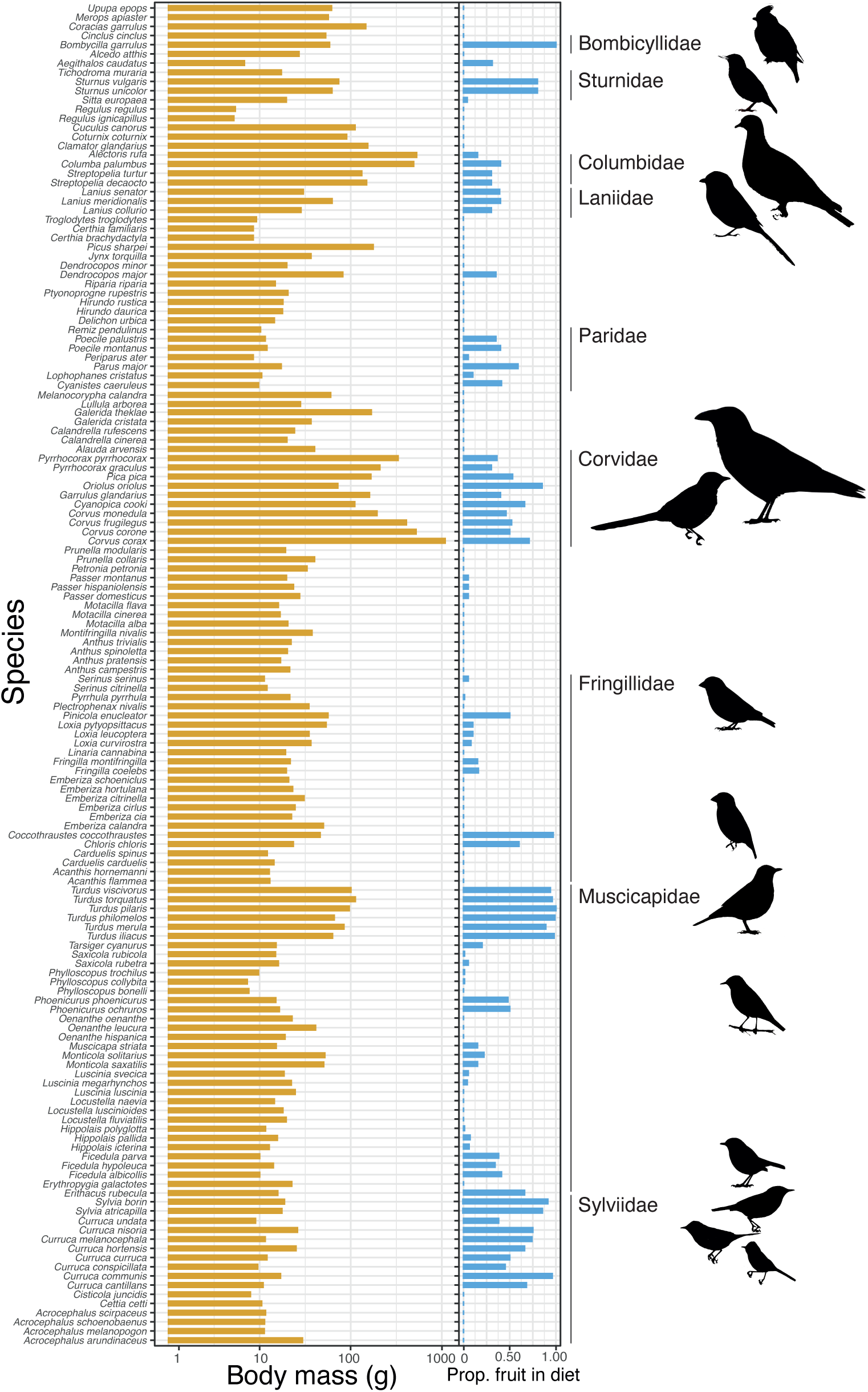
Body mass range variation (left, orange bars) and proportion of fruit in the diet (right, blue bars) for avian species of the Iberian Peninsula. Families including strongly frugivorous species are highlighted.

**Table 1:** Summary of the linear regression models for the percent of fruits in the diet (frupv) on avian species traits, separately for ecomorphological and digestive anatomy traits. masses are in g; linear measurements in mm. GPT, gut passage time (min); transit time (mm/min) estimated as intestine length/GPT. The whole model (all predictors included) is highly significant (*R*^2^ = 0.655*, F* = 4.48, *d.f.* = 11, 26, *P* = 0.00081). When analyzed separately, for ecomorphological traits: *R*^2^ = 0.307*, F* = 4.28, *d.f.* = 6, 58, *P* = 0.001; for digestive anatomy traits: *R*^2^ = 0.555*, F* = 6.45, *d.f.* = 6, 31, *P* = 0.0002. All variables *log*_10_-transformed prior to analysis.

| Trait | Estimate | Std. Error | $t$ | $P$ | | Relative contrib. $R^2$ |
| --- | --- | --- | --- | --- | --- | --- |
| *Ecomorphological traits* |  |  |  |  |  |  |
| Intercept | -0.471 | 0.821 | -0.57 | 0.570 | n.s. |  |
| Body mass | 0.209 | 0.301 | 0.69 | 0.493 | n.s. | 0.0986 |
| Wing length | -0.976 | 0.492 | -1.98 | 0.057 | ∼ | 0.0713 |
| Tail length | 0.711 | 0.352 | 2.02 | 0.053 | ∼ | 0.0657 |
| Tarsus length | 0.138 | 0.275 | 0.50 | 0.620 | n.s. | 0.0647 |
| Culmen length | -0.636 | 0.291 | -2.19 | 0.038 | * | 0.0543 |
| Gape width | 1.265 | 0.392 | 3.23 | 0.003 | ** | 0.1652 |
| *Digestive anatomy traits* |  |  |  |  |  |  |
| GPT | 55.932 | 31.491 | 1.78 | 0.087 | ∼ | 0.0449 |
| Gizzard mass | -0.434 | 0.110 | -3.93 | 0.001 | *** | 0.1772 |
| Liver mass | 0.109 | 0.180 | 0.61 | 0.555 | n.s. | 0.1046 |
| Intestine length | -55.834 | 31.447 | -1.78 | 0.088 | ∼ | 0.0526 |
| Transit time | 56.025 | 31.514 | 1.78 | 0.087 | ∼ | 0.0721 |

The avian assemblage of 146 species (Fig. 1) includes *n* = 56 (38.36%) seed disperser (SD) species, *n* = 6 (4.11%) pulp consumers (PC), *n* = 11 (7.53%) pulp consumer-seed disperser (PC/SD), *n* = 10 (6.85%) pulp consumers/seed predators (PC/SP), and *n* = 63 (43.15%) non–frugivorous (NF). Yet even for NF species sporadic records of frugivory exist (Turcek, 1961). SD species typically pick fruits and swallow the fruits whole after a short handling time (gulpers). PC and PC/SD species, in contrast, pick the fruit and handle it to tear-off pieces of pulp, in most cases discarding the seed(s). On occasions, PC species such as *F. coelebs*, *G. glandarius*, or *P. ater* and other parids leave the plants with fruits in the bill to handle in other nearby perches, just behaving as seed dispersers (PC/SD). Yet ample variation exists in the details of how fruits are handled; and handling behavior is not species-specific but varies depending on the fruit species the bird is consuming.

SD species typically swallow fruits whole (gulpers) after a short handling of the fruit in the bill. PC and PC/SD species in contrast, pick the fruit (finches, European jay) or peck it while attached to the peduncle (e.g., tits) to tear-off pulp. PC/SP species either pick the fruits to extract the seed contents after cracking the seeds in the bill (greenfinches, Hawfinch when foraging on one-seeded fruits) or extract the seeds while pecking the fruit still attached to the peduncle (greenfinches; when foraging on multi-seeded fruits, such as the cones of junipers), actually behaving as seed predators in both cases. Yet in some plant species they behave as pulp consumers and for this reason were grouped in PC/SP. These differences have a distinct effect on foraging rates and fruit handling efficiencies (see below), mostly implying lower fruit handling rates for PC and PC/SP species.

### 4.2 Ecomorphological correlates of frugivory of fruit consumption

As expected, there were significant differences among the five frugivory types in the reliance on fruit for food (*F* =46.1, *P <* 2*e^−^*^16^, *d.f.*= 4, 140), with SD and PC/SD showing higher dependence on fruits compared with the more occasional consumption by the rest of species. Across species the percentage of fruit in diet (*frupv*) is significantly and positively related to gape width, and negatively to culmen length (Table 1); this variation in *frupv* is also explained marginally by other traits, being positively associated to tail length, and negatively with overall body size (e.g., wing length) (Table 1, ecomorphological traits).

In relation to digestive system characteristics (Table 1, digestive anatomy traits), *frupv* decreased significantly with increasing gizzard mass and longer intestines, but increased significantly with higher food passage rate through the intestine (marginally with both *gpt* and transit time). Greater reliance on fruit food thus appears associated with rapid digestion times, with reduced digestion rates due to rapid food processing in longer intestines and smaller gizzards. In the multiple regression analysis (Table 1) the effect of body mass is probably obscured by its strong covariation with gizzard mass (*r* = 0.80473, *t* = 11.5, *P* =*<* 2*e^−^*^16^, *d.f.* = 72).

The set of traits (Table 1) explained a 65.47% of variation in the percentage of fruit in diet (*frupv*) across species. Taken together, ecomorphological variables (Table 1) explained a 47.51% of variation in the percentage of fruit in diet (*frupv*) across species, and the digestive anatomy traits accounted for 52.49%. Just body mass and gape width account for 26.4% of variation in *frupv*, and just gizzard mass and transit time account for 24.9%.

These effects on fruit foraging rates translate into variation in the per visit feeding rate (no. fruits consumed per unit time and number of seeds removed per visit to the plants): the number of fruits consumed per min significantly increased across species with body mass (*r* = 0.4701, *F* = 133, *P* =*<* 2*e^−^*^16^, *d.f.* = 1, 476; no. fruits per min= -0.520 + 0.555[body mass], on *log*_10_-transformed data) (Fig. 2), the regression using data of *N* = 478 feeding rates of individual birds foraging at different fruit species (Suppl. Mat. Table 4). This also resulted in higher ingestion rates of fruits per visit to plants with increasing body mass (see Suppl. Mat. Fig. 1 and Suppl. Mat. Table 4).

**Figure 2:**
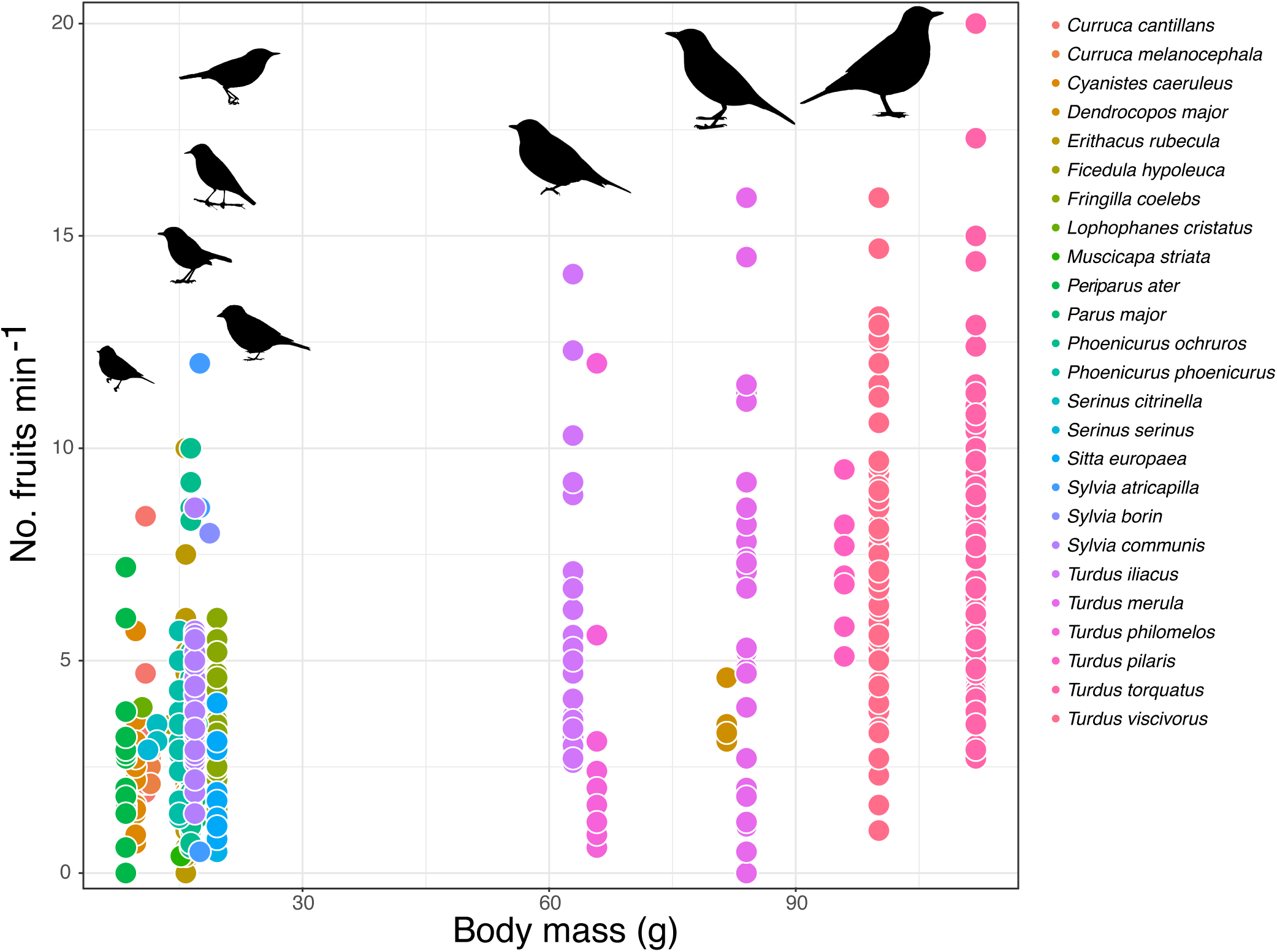
Relationship between body mass and per-visit fruit ingestion rate across Mediterranean avian frugivore species.

A canonical discriminant analysis (CDA) on both external traits and digestive characteristics of avian species (Fig. 3, Suppl. Mat. table 2) indicates distinct patterns of ecomorphological variation associated with fruit consumption. The CDA revealed highly significant differences between the groups of avian frugivores, discriminating between SD and PC-PC/SD species along CDA1 and both groups contrasting with PC/SP and NF species both along CDA1 and slightly along CDA2 (Table 2, Fig. 3). SD species are characterized by larger overall body size, wider gape, higher food passage rate (mm·min*^−^*^1^) through the digestive apparatus (due to a combination of shorter gpt in relatively shorter intestines), and larger liver. These characteristics markedly contrast with the longer gpt, more robust bills, larger gizzards and longer intestines of the PC and PC/SD species and the PC/SP group as well (Fig. 3). Non-frugivores are largely characterized by overall smaller body size and contrasting trends in the above variables (Fig. 3).

**Figure 3:**
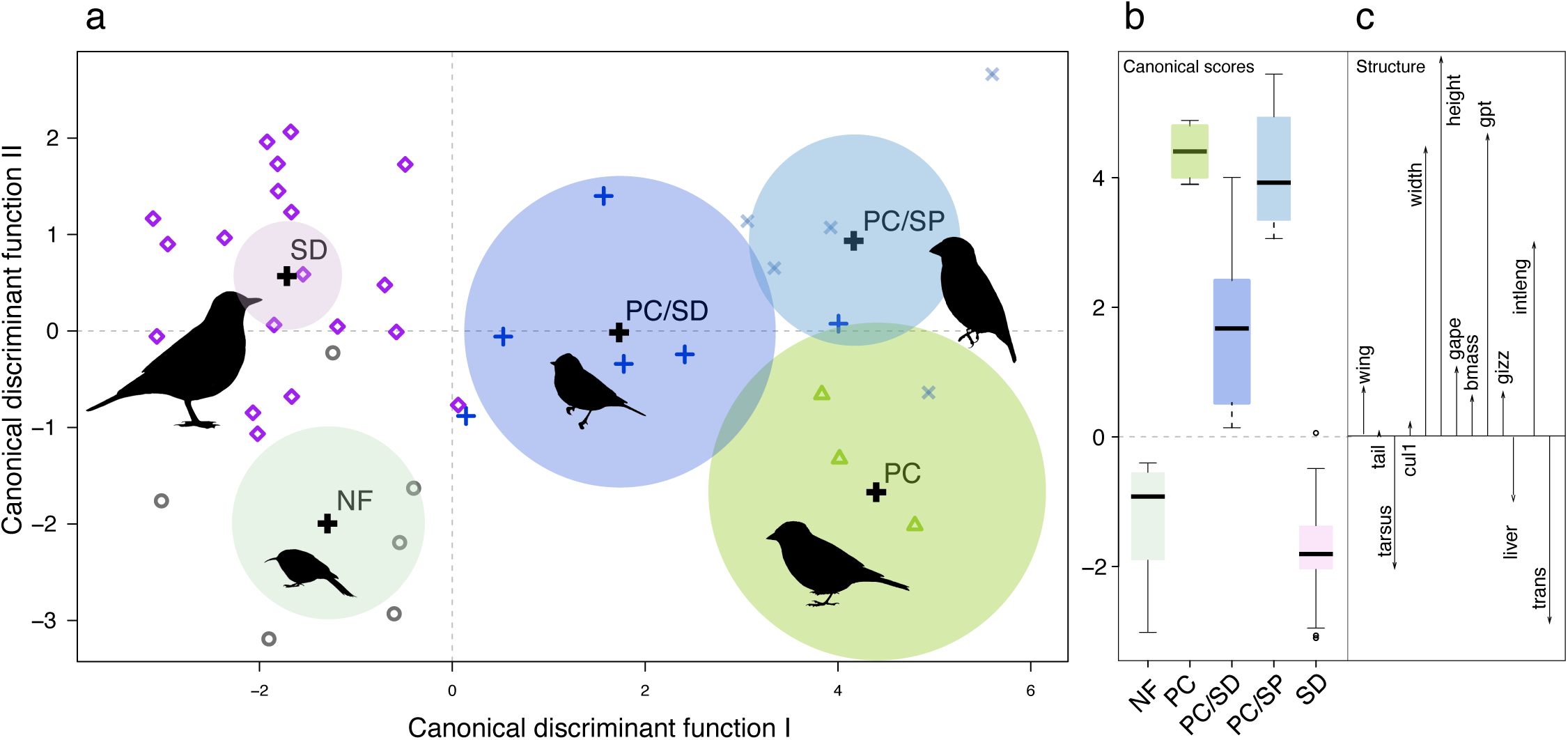
Results of a canonical discriminant analysis (CDA) based on morphological and digestive apparatus characteristics of Mediterranean bird species, grouped by type of frugivory (a). SD, seed disperser species; PC, pulp consumer species; PC/SD, pulp consumer/seed disperser species; PC/SP, pulp consumer/seed predator species, and NF, non-frugivorous species. Coloured circles indicate the 95% equiprobability areas for each group. (b) Mean canonical scores of each group along the first discriminant function; black horizontal lines denote the mean score, with the coloured rectangle encompassing ±1 SE and lines spanning the 10th and 90th percentiles (with outlier values denoted as small circles) Arrows (c) indicate loads of ecomorphological traits on the first discriminant function. Variable abbreviations as in Suppl. Mat. Table 2.

**Table 2:** Summary of the canonical discriminant analysis (CDA) of avian morphological and digestive apparatus traits between the major feeding groups in relation to fruit consumption: SD, seed disperser species; pulp consumer, pulp consumer/seed disperser, and seed predator (PC; with PC/SD, and SP, respectively) and non-frugivorous species (NF). The standardized coefficients for the traits on the first three discriminant functions are shown.

|  | D1 | D2 | D3 | D4 |
| --- | --- | --- | --- | --- |
| Canonical R<sup>2</sup> | 0.817 | 0.759 | 0.416 | 0.388 |
| Eigenvalue | 4.457 | 3.152 | 0.713 | 0.635 |
| Percent | 49.76 | 35.19 | 7.96 | 7.09 |
| Cumulative | 49.8 | 84.9 | 92.9 | 100.00 |
| *F* | 3.08 | 2.36 | 1.41 | 1.52 |
| *P* | 2.2·10<sup>-6</sup> *** | 0.0012 ** | n.s. | n.s. |
| Standardized coefficients |  |  |  |  |
| Wing length | -2.8722 | -2.1309 | -3.7700 | - |
| Tail length | 0.7178 | 1.2011 | 0.6578 | - |
| Tarsus length | 0.5812 | 1.8603 | -2.4320 | - |
| Culmen length | -1.2806 | 0.5442 | 0.3142 | - |
| Culmen width | 2.0139 | -1.3542 | -0.0164 | - |
| Culmen height | 0.5745 | -1.4673 | -2.0934 | - |
| Gape width | 0.8918 | 6.7227 | 2.9292 | - |
| Body mass | 3.0028 | -1.5491 | 0.0533 | - |
| Gut passage time | 0.7313 | 2.0246 | 1.1656 | - |
| Gizzard mass | -2.5885 | -3.5995 | 0.9901 | - |
| Liver mass | -1.0701 | -1.1459 | 0.492 | - |
| Intestine length | -0.3441 | -0.2868 | 1.6579 | - |
| Transit rate | 0.2013 | 0.0743 | 0.5934 | - |

### 4.3 Foraging for fruits

We combined data of feeding rates and fruit foraging behaviors for avian frugivores on 10 different plant species (Suppl. Mat. tables 3, 4, 5). A typical feeding sequence of a frugivorous bird foraging at a fruiting plant involves a sequence starting when entering the plant (visiting) until it leaves the plant after handling some fruits. When foraging on the ground for fallen fruits the bird enters the area of the canopy projection, where fallen fruits tend to accumulate. Once at the plant the bird moves or stays perched and starts a sequence of moves seeking and picking fruits. Once the fruit is picked it is handled with a variety of behaviors, possibly ending with swallowing the fruit or dropping it (or the seeds) beneath the parent plant. Picking the fruit may also fail, for example when foraging on nearly ripe fruits that are difficult to detach from the peduncle. We have never recorded a species with a frequency of fruit picking failures *>* 10% of the picking attempts (Suppl. Mat. table 5), yet except for *Turdus* species when foraging on species with limited accessibility. For example fruit picking failures for *T. torquatus* and *T. viscivorus* when feeding on *Lonicera arborea* are 17.3 and 28.9%, respectively; while fruit picking failures *T. viscivorus* when feeding on *Taxus baccata* amount to 19.8%. These thrushes frequently loss their position while foraging on *L. arborea* due to the thin ramets that hold the infructescences, that frequently broke due the weight of the birds. The thin branches bend when the birds forage, forcing them doing frequent wing flaps, losing equilibrium and finally breaking the branches and leaving the plant with the feeding sequence interrupted.

Birds used seven basic feeding maneouvres while foraging on fruits (Suppl. Mat. tables 3, 4; Suppl. Mat. Fig. 2). Most SD and PC species glean for fruits from the branches using quick moves and picking the fruits. Using a nearby branch as a perch (most frequently the branch subtending the infructescence from which the fruit is picked), the fruits can be picked either upwards, with the bird elongating the body to reach a fruit high from its position. Most times however the picked fruits are from infructescences just in front of the bird. Finally, some species may obtain fruits by picking them from a position well below the branch used to perch. The bird literally hangs downwards from the branch, picks the fruit and regains its position to handle and swallow the fruit. Only Blackcaps, Whitethroats and Redwings, Blackbirds and Mistle thrushes were observed picking fruits downwards and being able to regain their perched positions. Birds foraging for fruits from the branch quite often loss equilibrium, flap the wings, correct their perching position or move to another perch. This is clearly seen in the larger species (e.g., thrushes, pigeon, Azure-winged magpie, Jay). Some species, e.g., Robin, flycatchers, stonechats, redstarts, use predominantly on the wing captures to pick the fruits or sallies from a perch to pick the fruit on the pass and either get back to the perch or go to another nearby perch (Suppl. Mat. Fig. 2). On the flight sallying is also used by thrushes, especially when foraging for fruits later in the season when most fruits available are located at the distal parts of branches or infructescences. While sallying typically does not involve stalling in flight to pick the fruit, the redstarts, flycatchers, Robin and stonechats frequently flycatch the fruits from a perched position: they perch for a relatively long period of time and suddenly make a quick jump to pick a fruit while momentarily flying stalling in front of it to immediately return to the same perch or a different one (Suppl. Mat. Fig. 2). Finally, some species (e.g., Robin, redstarts) may sally to the ground to pick fallen fruits, while others (e.g., thrushes, Robin) pick fallen fruits on the ground.

The PC and PC/SD as well as the PC/SP species have characteristic modes of fruit foraging and handling (Suppl. Mat. Fig. 2). The fruit is pecked without detaching from its peduncle, obtaining pieces of pulp that are ingested, usually from a branch. The seeds may remain attached to the infructescences with the pulp pecked (e.g., in *Taxus*, *Rubus*, *Prunus*) or just the fruit open with pieces of pulp and content missing (as e.g., in *Arbutus*, *Juniperus*, *Pyrus*, *Malus*, some *Sorbus*). Some species (e.g. Jay, finches) are able to handle the fruit in the bill while tearing off the pulp to ingest it. Yet most species picking the fruits to consume the pulp (e.g., tits), peck directly the fruit or use an assisted feeding by holding the fruit in their feet against the branch and then pecking the pulp; the seed(s) are dropped to the ground. Finally, PC/SP species (e.g., Hawfinch, Greenfinch) either peck the fruits without detaching them (e.g., in *Juniperus phoenicea*, *Taxus*, *Prunus*) or picking the fruit (e.g., in *Pistacia*, *Prunus*, *Taxus*, *Crataegus*) and extract the seeds to crack them open in the bill and ingest the seed content. This behavior may vary (either picking or pecking) for the same plant species, with the birds behaving at times as PC species feeding on just the pecked pulp.

The immediate result of this diversity of foraging modes is ample variation in feeding rates and foraging speed. Suppl. Mat. Fig. 3 illustrates this variation by means of step graphs (Cody, 1968). The slope of the step graph is proportional to the foraging speed (no. fruits handled per min). Some species, especially the SD species, are very quick foragers, almost swallowing fruits immediately after they pick them (e.g., redwings and mistle thrushes, blackcaps, garden warblers and whitethroats). This frequently involves very rapid manoeuvres while gleaning during relatively short visits to the plants. The foraging speed of these small and medium-sized SD species is notably higher than the speed of larger SD species (corvids, pigeon). These SD species characteristically have very short search times for fruits and extremely short handling times. Fruits are swallowed as they are picked, although sometimes birds take longer handling times while repositioning the fruits in the bill before swallowing. This is clearly seen when frugivores forage on fruits whose size is large in relation to bill size and gape width. If the fruits are too large then handling times typically are longer. In the step graphs (Suppl. Mat. Fig. 3) this is reflected in longer inclined segments for each fruit (each step), as the length of these segments is proportional to fruit handling time. In contrast, PC and PC/SD (and also PC/SP) species have notably longer step segments, especially in the inclined portion of the step (Suppl. Mat. Fig. 3): tits and finches have long handling times when they pick the fruits and hold them in the bill while tearing-off pulp; then they perform very quick moves to pick other fruit and stop again with a long handling time. The same occurs in nuthatches and Jay, that pick fruits and handle them with the foot, holding the fruit against the perch branch while they peck and tear-off pulp pieces. Nuthatches also pick fruits to a nearby large branch where they place the fruit in a crevice of the trunk and then peck the pulp and/or seeds, cracking them open. Obviously these feeding modes involve long handling times in these PC, PC/SD and PC/SP species (Suppl. Mat. Fig. 3). Finally, birds using flycatching and sallying manoeuvres typically have long horizontal segments in the step graphs (Suppl. Mat. Fig. 3) illustrating their long perching time, and very short and steeper vertical segments of the steps indicating the short and quick hops to flycatch the fruits. These are typical manoeuvres of robins, redstarts, flycatchers, and stonechats.

### 4.4 Fruit handling

Most SD species, especially the largest-bodied species, are gulpers that swallow fruits whole, frequently with very high handling success (¿95% of the handled fruits swallowed; Suppl. Mat. Table 5). Across species, a positive correlation exists between % fruits swallowed and body mass (*r* = 0.6573*, P <* 0.005; Suppl. Mat. Table 3) suggesting increasing constraints on fruit handling for smaller species. In contrast, PC and PC/SD species typically handle the fruit externally and do not ingest, frequently dropping the fruits or seeds. Some species may change fruit handling depending on fruit species; for example, nuthatches pick *P. mahaleb* fruits and use assisted feeding while pecking the pulp on fruits/seeds attached to bark crevices, but they swallow fruit whole when feeding on *T. baccata*.

Failures to pick the fruits while foraging are infrequent (*<* 10%; Suppl. Mat. Table 5), and mostly involve instances where the birds are trying to pick unripe fruits that are still tightly attached to the peduncles. Also, small birds like *P. ater* or those that use hovering or sallying (*Phoenicurus* spp.) show infrequent failures of this type, most likely associated to the lower precision of on-flight foraging manoeuvres. More common are fruit handling failures when holding a picked fruit in the bill just before ingestion. In this position, the fruit is often repositioned in the bill prior to swallowing and may fall to the ground, both in large and small species. Yet, there is no negative correlation between gape width and % failures and dropping of fruits across the individual observations of feeding rates (*r* = −0.0363*, P* = 0.38*, d.f.* = 575). Failures to handle fruits appear more related to misfits between gape width and fruit size, with a negative correlation between these two variables (*r* = −0.11025*, P* = 0.007*, d.f.* = 581)(Suppl. Mat. Table 5) suggesting that larger differences between fruit size and gape width of the foraging species increase the chances of foraging failures.

Increasing trait matching between fruit size and bird size appears an effect of overall body size and the correlation between body mass and gape width, favouring more efficient feeding rates for larger species able to handle efficiently a wider range of fruit sizes (Suppl. Mat. Fig. 2). The frequency distribution of fruit diameter for plant species of the Iberian Peninsula (Fig. 4) suggests that birds with gape width *<* 7*mm* (modal fruit size diameter for *n* = 111 fruit species; Suppl. Mat. dataset) may have handling constraints to efficiently consume a large fraction of larger-sized fruit species. From the seed dispersal perspective the lack of fit between fruit size and gape width causes significantly increased proportion of fruit and seeds being dropped beneath the plants (*r* = −0.40416*, P* = 2.3*e^−^*^16^*, d.f.* = 581 for the relationship between size.fit and pct.beneath)(Suppl. Mat. Table 5).

**Figure 4:**
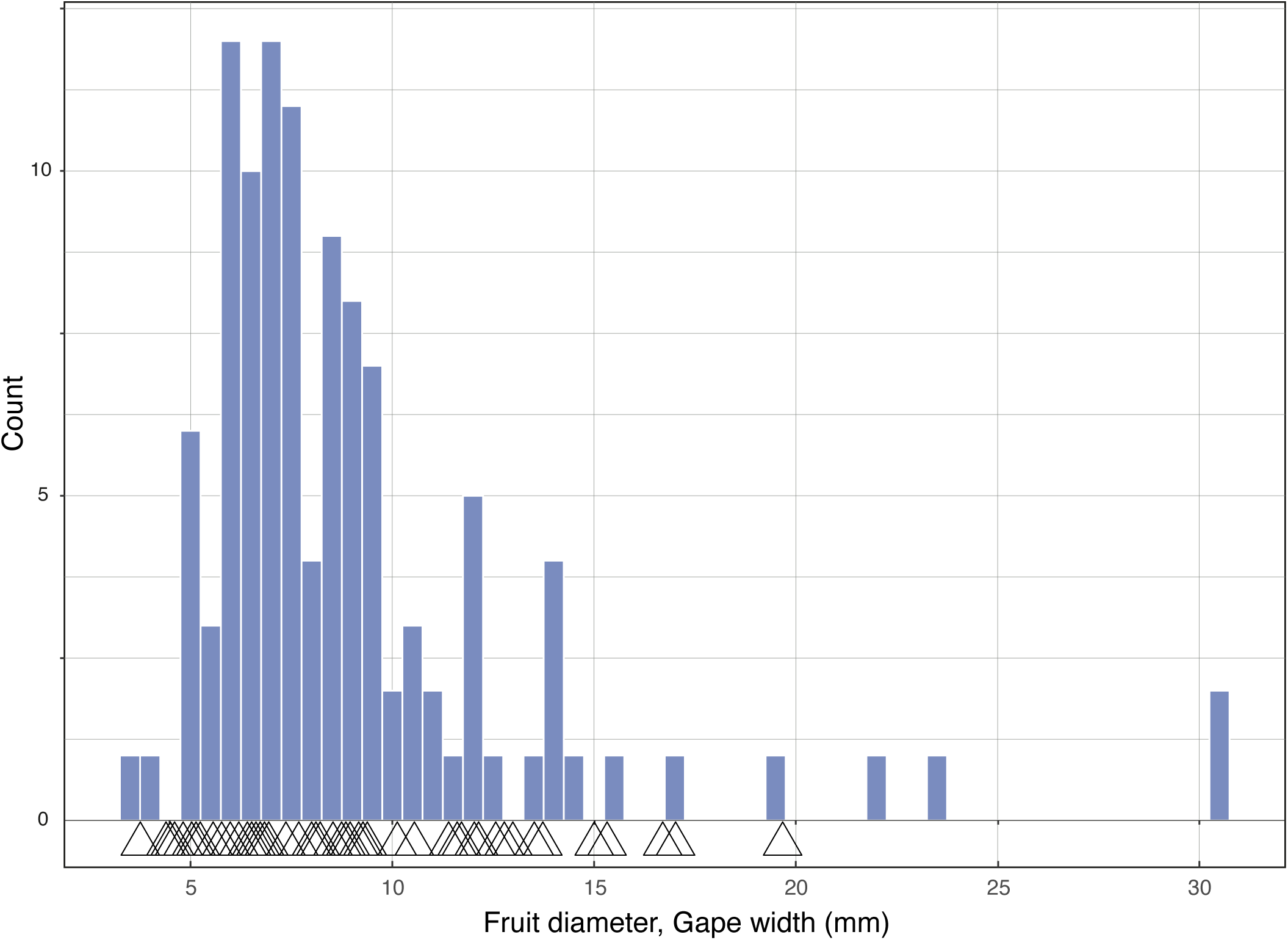
Frequency distribution of fruit diameter (mm) for Mediterranean scrubland fleshy fruited plant species (bars) (see Herrera, 1984b; Valido et al., 2011; Jordano, 2014). Mean gape widths (mm) of avian species are indicated by small triangles along the abscissa.

## 5 Discussion

The avian species assemblage studied includes most of the Iberian Peninsula species that may eventually consume fleshy fruits, excluding most non-passerine species like waders, ducks, seabirds, raptors, cranes, flamingoes, etc. that very sporadically may feed on fleshy fruits (Turcek, 1961). The species included illustrate an ample gradient of reliance on fleshy fruits and include some strong frugivores like the larger *Sylvia*, *Curruca*, and all the *Turdus* species (Jordano, 1988, and references therein). This frugivory gradient between fully insectivorous and mostly frugivorous species is paralleled by distinct ecomorphological traits, a characteristic feature that has been reported previously (e.g., Herrera, 1984b,a; Jordano, 1987a; Snow and Snow, 1988; Stiebel and Bairlein, 2008; Pizo et al., 2021; Carlo et al., 2022; Gomes et al., 2022). From this extreme of strong reliance on fruit food we found many species that rely on fleshy fruits to variable degree, from greater relevance in the diet (e.g., redstarts) to frequent consumption during migration (e.g., flycatchers, stonechats) to sporadic consumption especially in winter (e.g., *Phylloscopus* warblers). Larger species tend to consume fruits more frequently, and the proportion of diet made up by fleshy fruits is strongly and positively correlated with body mass and gape width, a trend that is maintained even within closely-related species (Jordano, 1987a; Stiebel and Bairlein, 2008). For example, the Sylvia-Curruca genera exhibit this gradient of variable reliance on fruits food (Jordano, 1987a), from the mostly frugivorous larger species (e.g., *S. atricapilla, S. borin*) to the smaller species (*C. conspicillata, C. undata*) that only occasionally feed on fruits, a trend replicated in other Muscicapidae (Jordano, 1987b). Such a gradient is also observed in other avian frugivore groups (e.g., Pizo et al., 2019, 2021; Carlo et al., 2022) and involves two main aspects among the most frugivorous species (Jordano, 1988): 1) a large fraction of individuals in a local population use fleshy-fruit food (i.e., use of fruit resources is extensive, despite variation among individuals) and 2) fleshy-fruits make up a large fraction of diet (i.e., *>* 90%, by volume).

### 5.1 Ecomorphology of fruit feeding

Ecomorphology constrains feeding behavior, search movements, feeding rates, and handling times, among other aspects of frugivores feeding ecology (Moermond and Denslow, 1985; Pizo et al., 2019). In the case of avian frugivores this severely limits many aspects of fruit feeding, given the diversity of fruiting displays and accessibility exhibited by fruiting plants. Fleshy fruits are particulate food that needs to be picked, handled, and eventually ingested adequately (Jordano, 2014); therefore, size constraints have been a major limiting factor in fruit use repeatedly reported (Wheelwright, 1985; Pratt and Stiles, 1985; Levey, 1987; Rey et al., 1997; Githiru et al., 2002; Pizo et al., 2019; Carlo et al., 2022). The nutritional characteristics of fleshy fruits impose rapid handling times and speedy feeding behaviors to foraging birds in order to overcome digestive bottlenecks created by the nutritional value and marked imbalance of nutrients in the fruit pulp (Karasov and Levey, 1990; Rio and Karasov, 1990; Jordano, 2014), frequently imposing tradeoffs and physiological constraints (Pizo et al., 2019) to the processing of fleshy-fruits, from handling to ingestion and digestion. Fleshy fruit profitability strongly depends on rapid processing and quick digestion and this is one of the main reasons why most avian frugivores exhibit mixed diets (e.g., fleshy fruits and arthropods) (Levey and Karasov, 1989). Mixed diets, on the other hand, require more generalized feeding modes and foraging behaviors, given that foragers with mixed diets often switch from one food type to the other even during short feeding bouts (Jordano, 1988; Pizo et al., 2019). Our study revealed that strongly frugivorous species show a distinct combination of body traits making this possible: larger body sizes, wide gape widths, relatively short intestines with rapid transit rates of food and relatively large livers. All these characteristics have been previously discussed in reference to local avian assemblages (Herrera, 1984b; Stiebel and Bairlein, 2008; Carlo et al., 2022) or specific frugivore groups (e.g., Jordano, 1984; Moermond and Denslow, 1985; Jordano, 1987a, 1988; González-Varo and Traveset, 2016). While large body mass determines greater amount of fruit that can be ingested during short visits to the plants, wider gapes determine accessibility to a wider range of fruit sizes and thus fruit species (Wheelwright, 1985; Forget et al., 2007). It is well documented that largely frugivorous diets require the ability to consume a diverse array of fruit species in order to complement nutritional imbalances (e.g., Jordano, 1988, 2014, and references therein). Across the Mediterranean species studied, the combination of larger body mass and reduced gizzard size, together with rapid food transit times relate to increased reliance on fleshy–fruit food.

### 5.2 Foraging for fruits

Foraging for fruit by avian frugivores includes a series of steps that can be organized hierarchically according to Sallabanks (1993) in relation to the scale of foraging “decisions” by the birds: 1) patch or area where to seek for fruits; 2) specific plant to visit; and 3) within-plant selection of individuals fruits. Our analysis refers mostly to step #3, regarding how birds are constrained by ecomorphology and this results in variation in feeding rates and hence, variation in intensity of fruit use. Foraging rates of avian frugivores within the plants are typically very short, especially when considering non-tropical frugivores (e.g., Snow and Snow, 1988; Stiebel and Bairlein, 2008) and compared to the longer fruit-feeding visits made by large tropical frugivores (e.g., Pratt and Stiles, 1983; Carlo et al., 2022).

The main contrast among species related to foraging rate on fruits concerns differences between gulpers (*SD*), swallowing fruits whole, and pulp consumers and seed predators (that infrequently may disperse seeds) (*PC*, *PC/SP* species). The former (e.g., *Sylvia*, some *Curruca*, *Turdus*) consistently show very rapid fruit handling and high ingestion rates, while the latter (e.g., finches, tits) are characterized by much longer fruit handling times and, consequently, lower handling rate. It is important to note that the covariation between number of fruits handled and time (feeding rate) varies among plant species, so that even *SD* species may have lower feeding rates when foraging on large-fruits species (e.g., *Turdus* spp. feeding on *Crataegus monogyna*) (Guitián et al., 2000) when compared to small-fruited species (e.g., *Pistacia lentiscus*) (Quintero et al., 2023) or those with aggregated fruits (e.g., *Rubus ulmifolius*) (Jordano, 1982).

### 5.3 Ecological correlates of frugivory

This paper examines the patterns of fruit use as a resource for a range of avian species. We have explored diverse traits, both ecomorphological and physiological potentially correlating with frugivory intensity, estimated as the % of total diet volume made up by fleshy fruits. The two groups pof traits contribute similar proportions of variance in *frupv* across species, with just body mass, gape width, gizzard mass, and transit time accounting together for 51.32% of the *R*^2^ between variables. These variables have been consistently identified as key variables influencing fruit use by avian frugivores (see e.g. Jordano, 1987a; Snow and Snow, 1988; Jordano, 2014; Pizo et al., 2019; Carlo et al., 2022).

### 5.4 Conclusion

Avian frugivores are widely diversified in the Iberian Peninsula, ranging from strong, almost exclusive fruit feeders (at least during migration and wintering) to species sporadically using the fruits as food. Major frugivorous groups include thrushes and *Sylvia* and *Curruca* warblers as well as a few muscicapids. For these species fleshy fruits not only made up a major fraction of the diet, but are also consumed by a large proportion of the individuals in a given local population. This highlights not only the relevance of fleshy fruits as a major food resource for many species, but also the important ecological role of these birds as seed dispersers for the plants (Jordano, 2014).

Frugivorous species may in turn add up a large fraction of the diversity and biomass of local Mediterranean avian communities, especially during autumn passage migration and wintering periods, this being a characteristic feature of these habitats and closely associated to the evolution of the Palaearctic-African bird migration system. The role of southern Spanish scrublands and forest as migratory passage and wintering grounds for European passerines cannot be understood without consideration of the fleshy fruit food resources they rely upon during these critical periods of their life cycles. The ecomorphological and foraging modes diversity described in this paper illustrate how fruit food use may have determined the evolutionary radiation of the Iberian avifauna, with adaptations ranging from distinct digestive anatomy details to combinations of bill and body traits. The reciprocal effect, the role of these frugivores in providing key ecological services for forest regeneration has not been sufficiently highlighted (Sekercioglu et al., 2016); this relevance is reflected also in their role in providing quick regeneration after fires on in degraded habitats (Mendoza et al., 2009; Garcia et al., 2010), and increasing plant diversity in monospecific plantations (Zamora et al., 2010). These effects are noteworthy, as in these diversified Mediterranean avian assemblages, just a relatively minor % of species relies extensively on fruit food *and* functionally disperses seeds undamaged.

## 6 Acknowledgements

This work is the result of many years of research and observation of frugivorous birds. We appreciate the help in the field and extensive discussions and ideas of Cristina García, Eugene W. Schupp, Myriam Márquez, Alfredo Valido, Manolo Carrión, JuanLu García-Castaño, Jesús G.P. Rodríguez, Jordi Bascompte, Francisco Rodríguez-Sánchez, Blanca Arroyo-Correa, Arndt Hampe, JuanPe González-Varo, Irene Mendoza and many others. Field work with the camera traps was assisted by Blanca Arroyo, Pablo Villalva, Pablo Homet, and Gemma Calvo.

Over the years our field work has been generously supported by the facilities at Parque Natural de las Sierras de Cazorla (Junta de Andalucía), Segura y Las Villas, ICTS-Reserva Biológica de Doñana (CSIC) and the administration of the Doñana National Park.

Completion of this manuscript was funded by the Spanish Research Agency (grants CGL2017-82847-P and PID2022-136812NB-I00 from the Agencia Estatal de Investigación), the Plan Propio de Investigación y Transferencia, University of Sevilla (2021-2022), and a LifeWatch ERIC-SUMHAL project (LIFEWATCH-2019-09-CSIC-13) with FEDER-EU530 funding.

PJ dedicates this work to the memory of Manolo Carrión and JuanLu García-Castaño for the many hours of field work and excellent funny moments shared with them.

## 7 Data archiving

Data and code for this paper are deposited in the GitHub repository https://github.com/PJordano-Lab/MS_avian_frugivores.

## Supplementary Material

### 1 Supplementary Material Tables

**Table 1: Suppl. Mat. Table S1.**
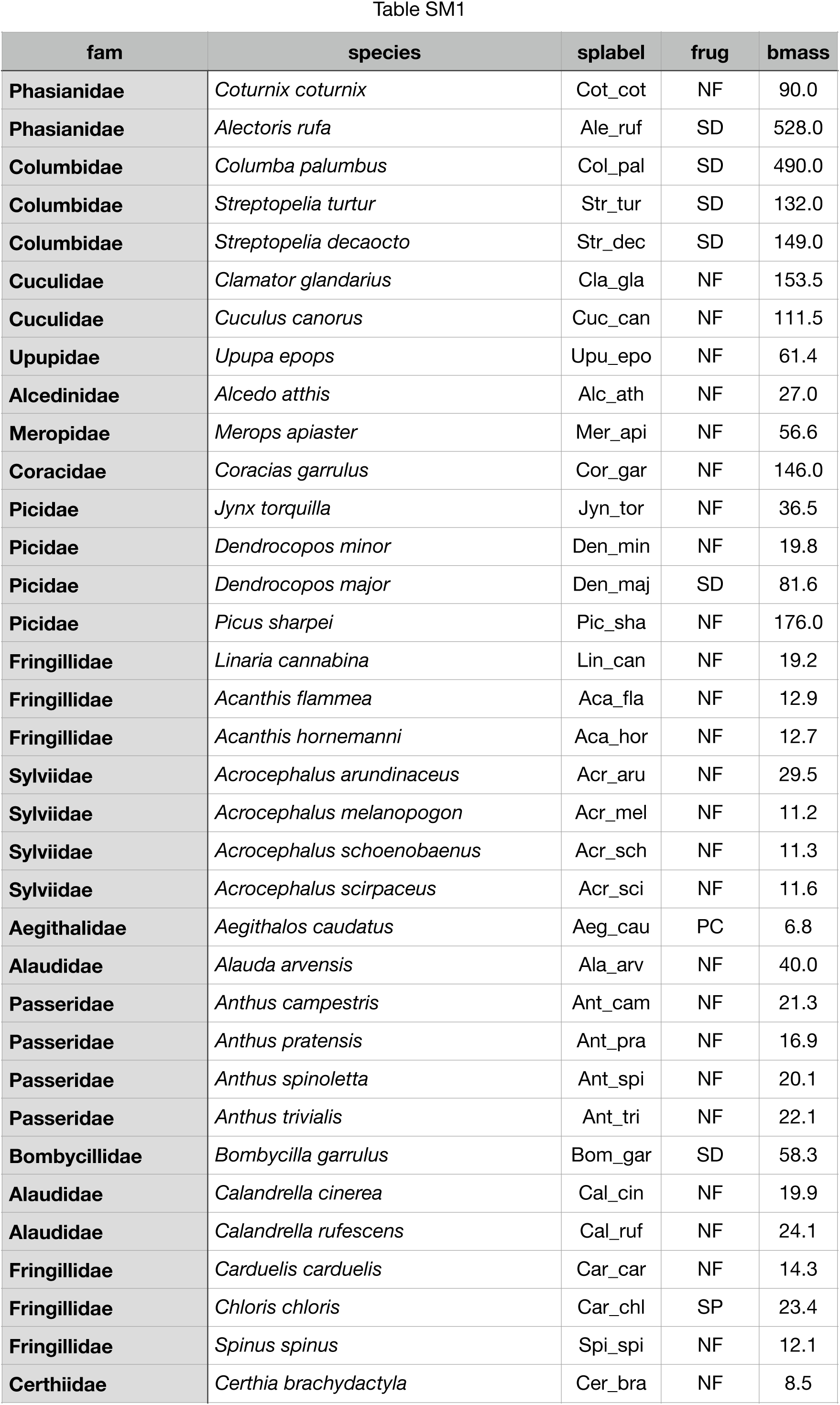
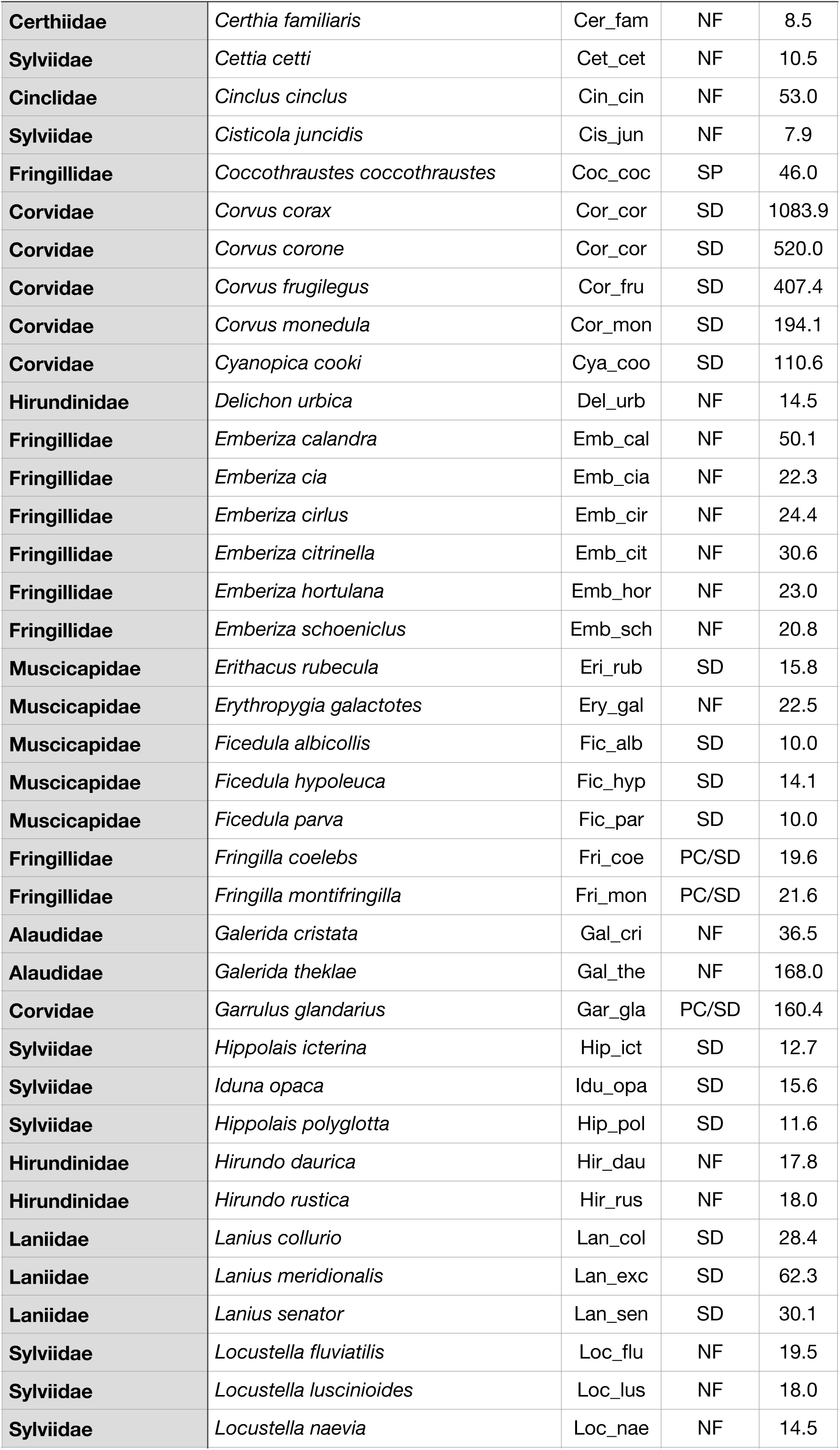
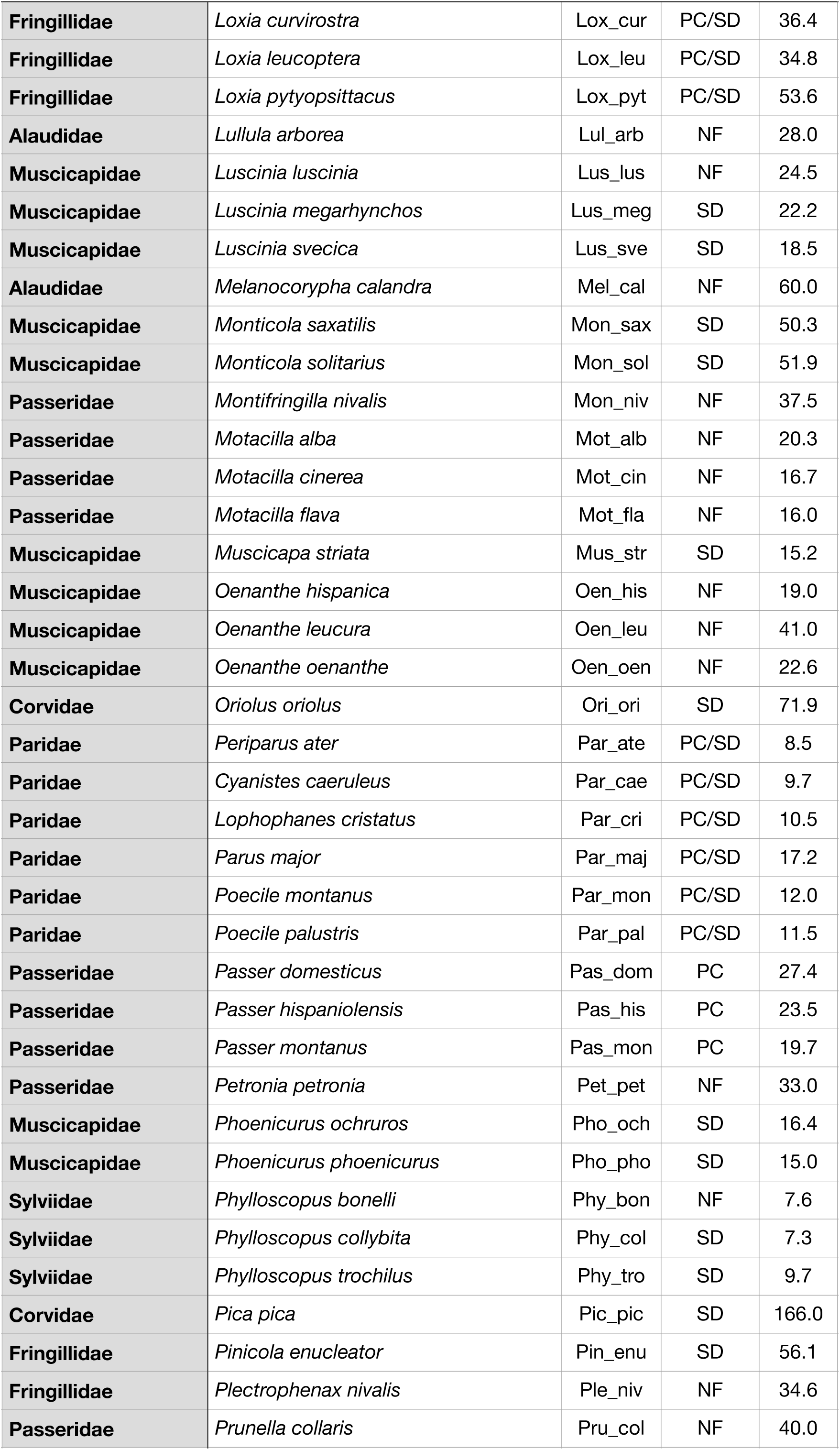
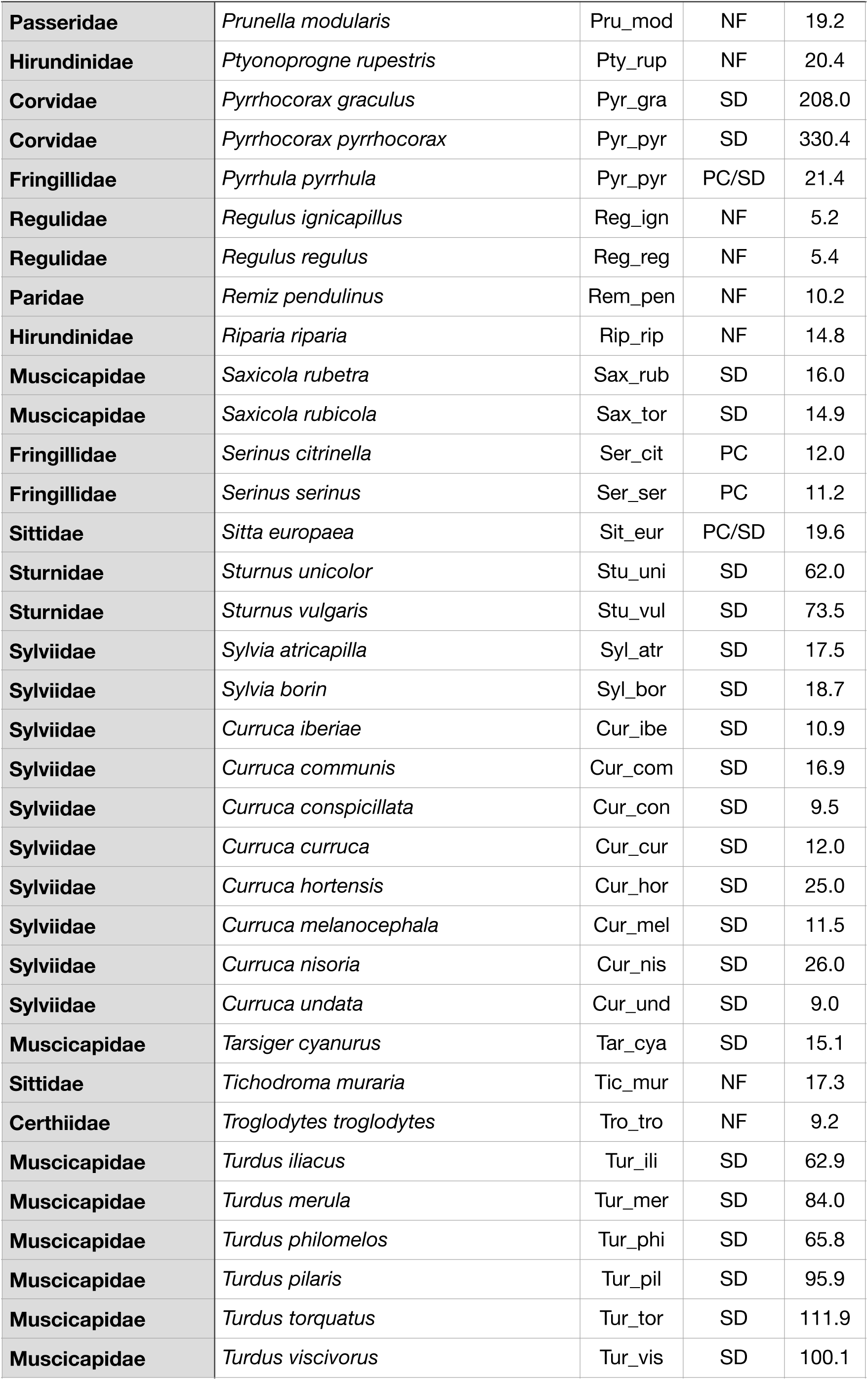
Avian species included in this study, types of fruit foraging, and body mass (g). Seed dispersers, SD; pulp consumers, PC; pulp consumer-seed dispersers PC/SD; seed predators, SP; and non frugivores, NF.

**Table 2: Suppl. Mat. Table S2.**
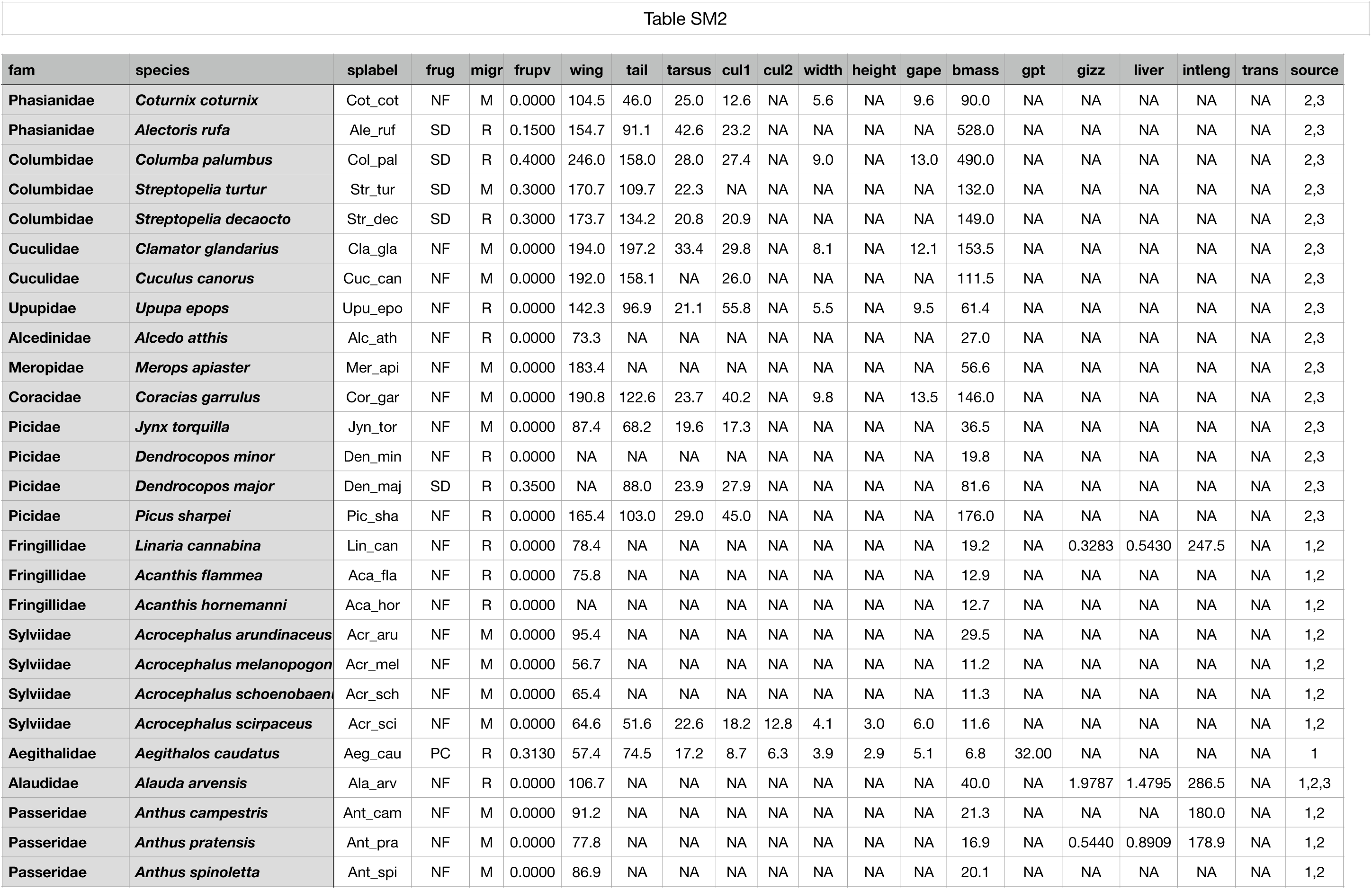
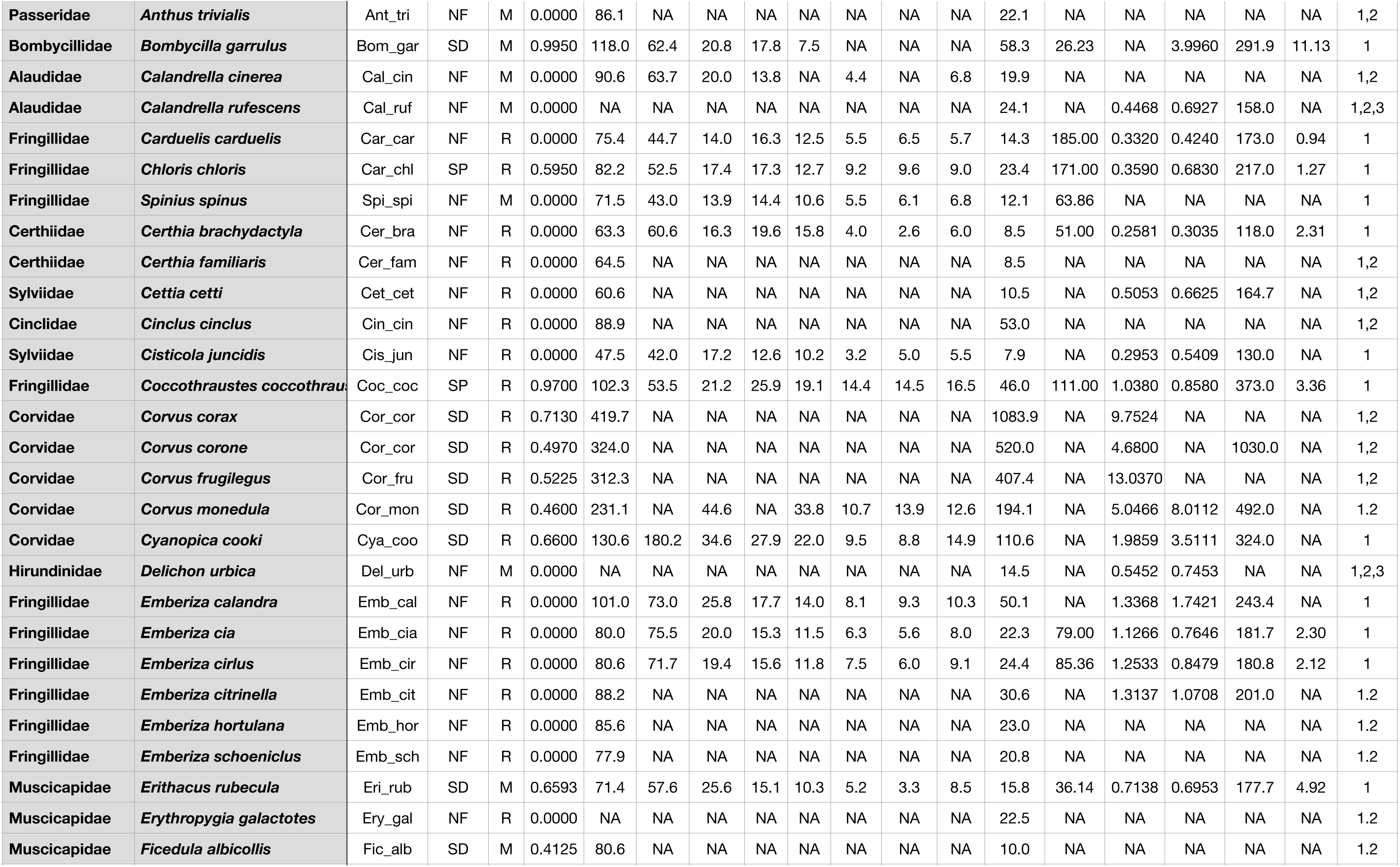
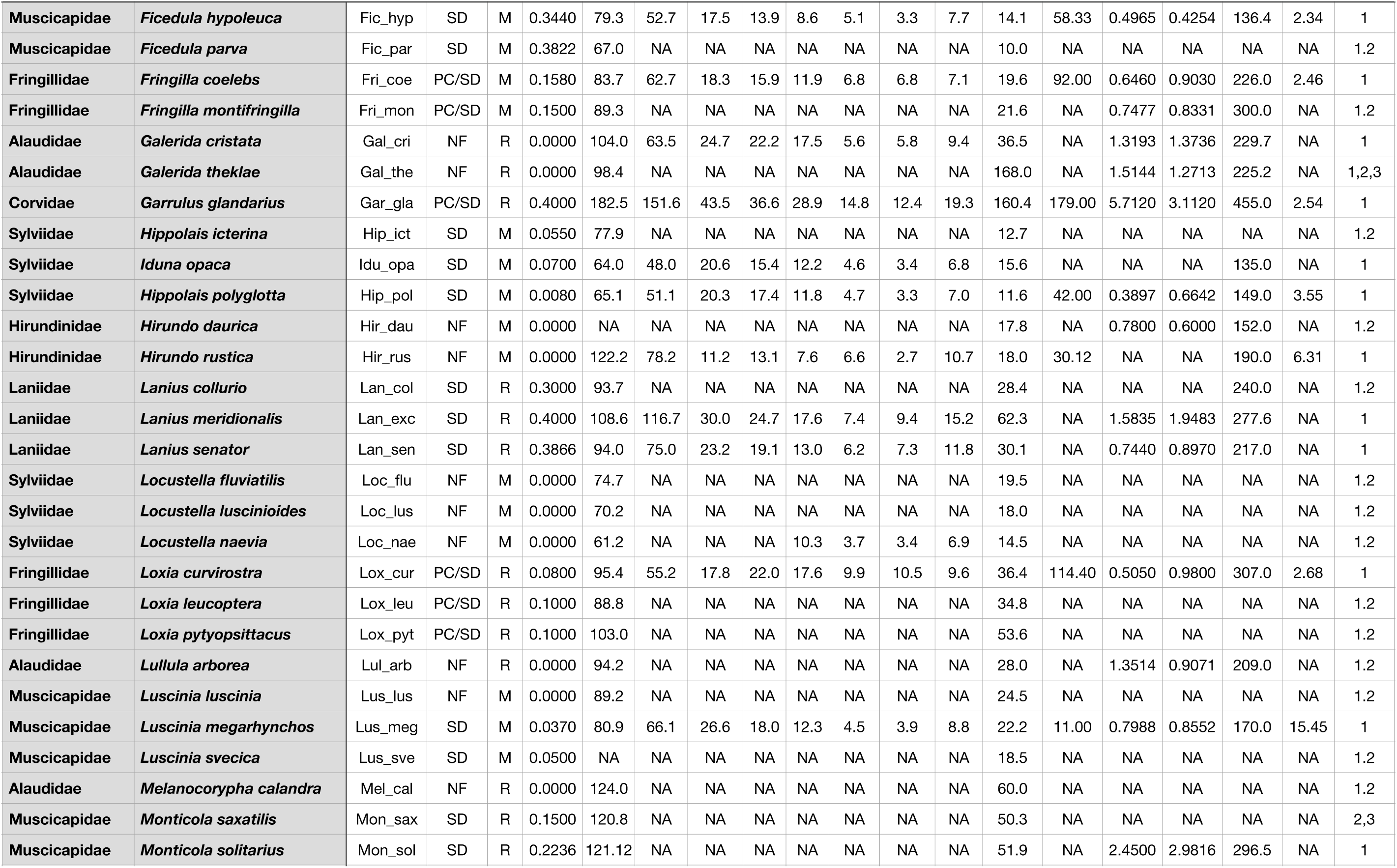
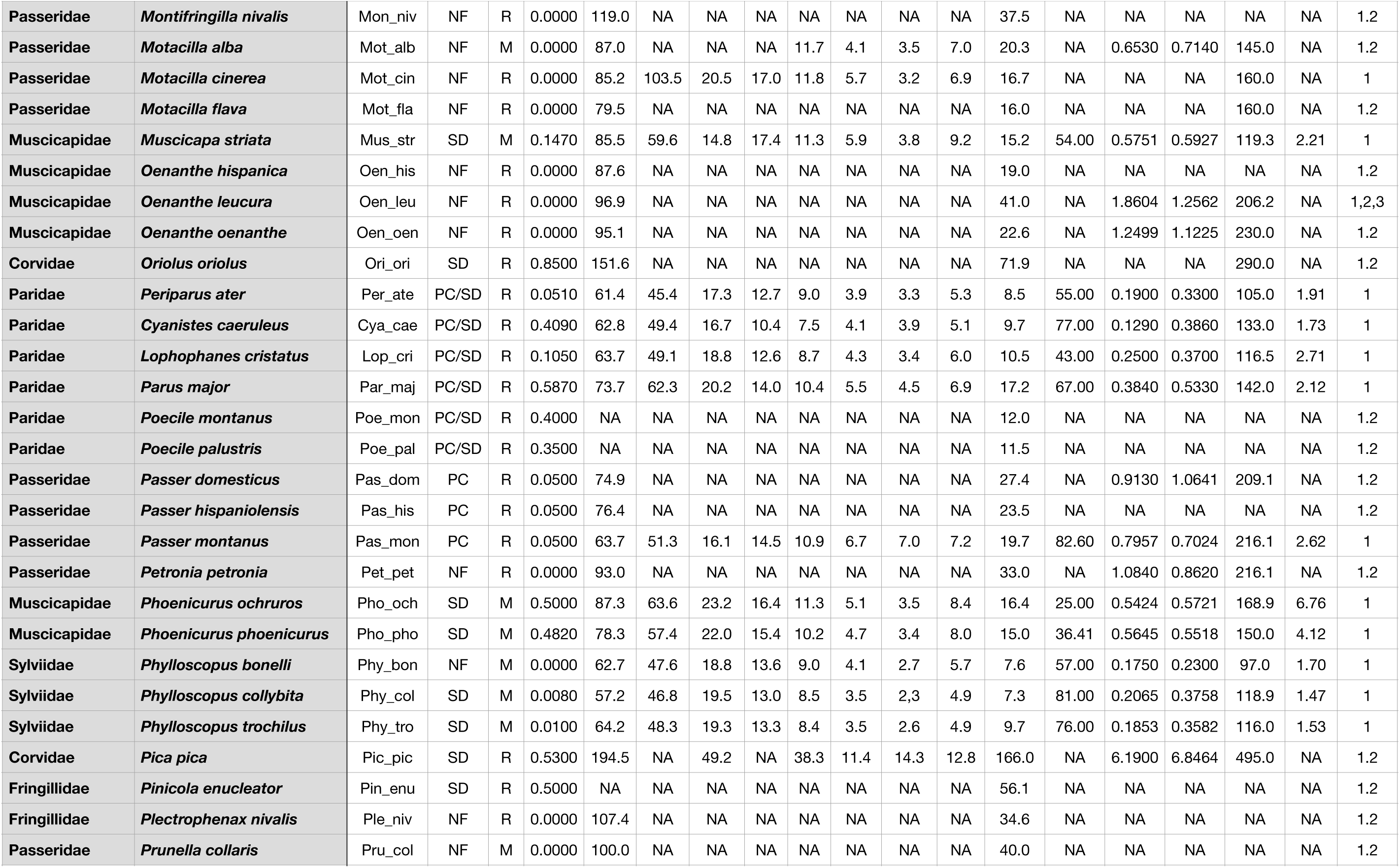
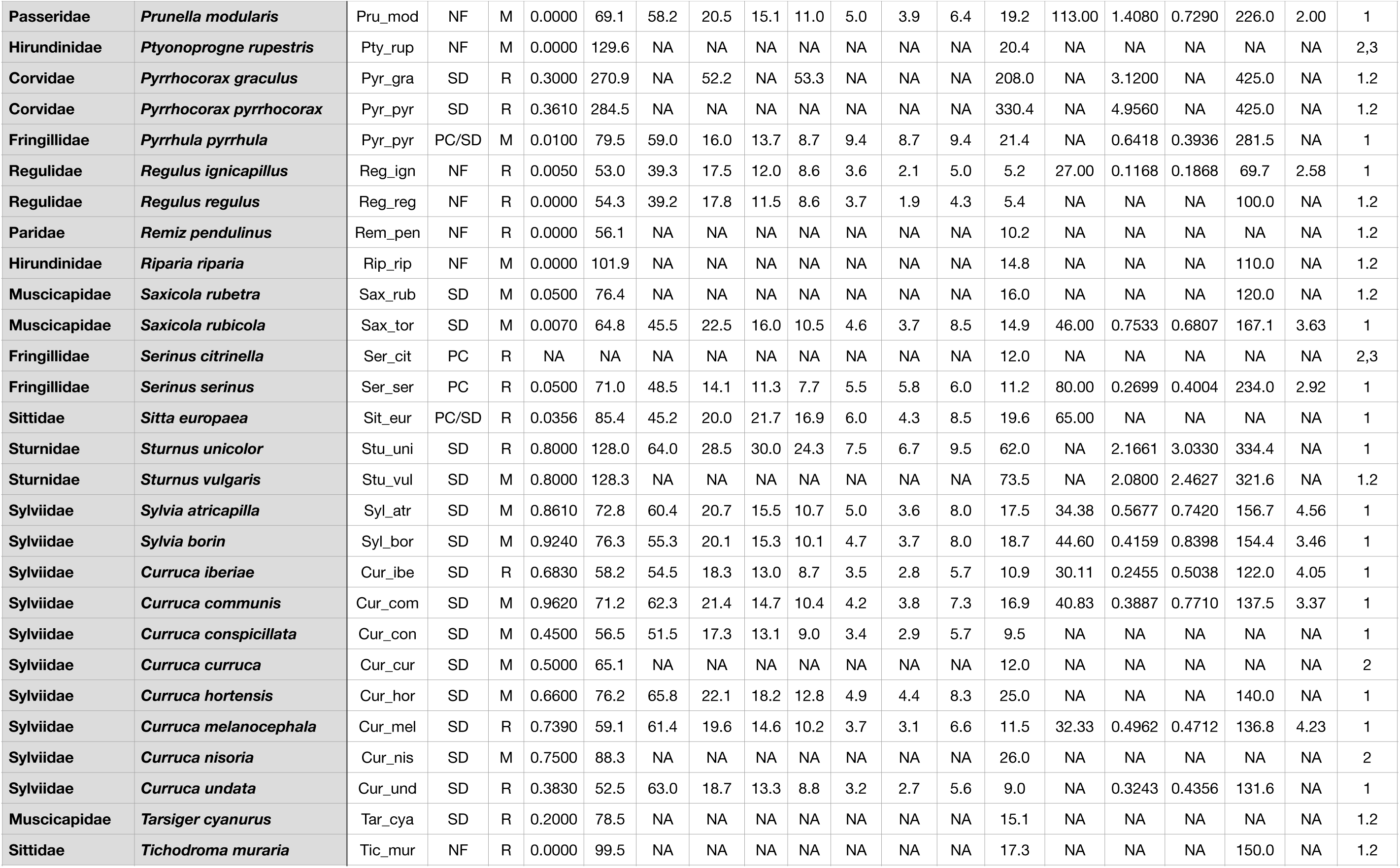
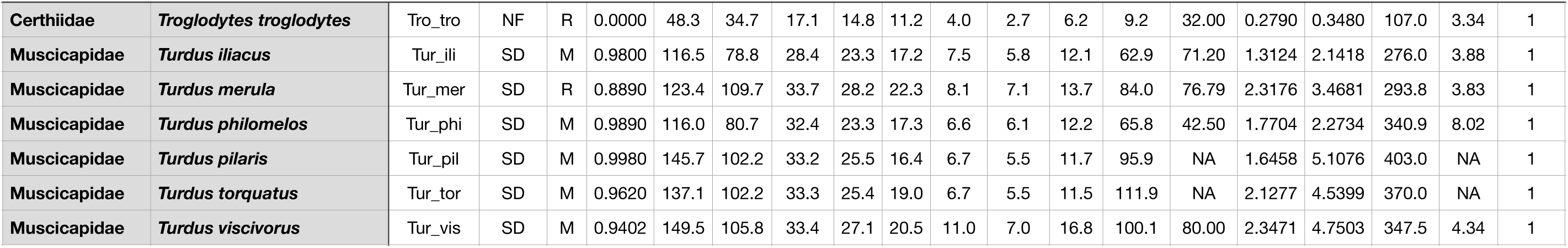
Frugivory types (frug), migratory status in S Spain localities (migr), proportion of diet volume made up by fleshy fruits (frupv), and ecomorphological characteristics of avian species included in this study. Traits are mean values and include: frug, type of frugivory: NF, non-frugivore; SD, legitimate seed disperser, usually swallowing whole fruit and dispersing seeds; PC, pulp consumer, usually tearing-off pieces of pulp and dropping seeds; PC/SD, pulp consumer-seed disperser, a PC species that occasionally ingests seeds and/or carries seeds away the plant without dropping them (included as PC); SP, seed predator. migr, migratory status in S Spain sites (dominant): R, resident (whole year); M, migrant. frupv, proportion of diet volume contributed by fleshy fruits. wing, wing length (mm), obtained from folded wing, stretched. tail, tail length (mm). tarsus, tarsus length (mm). cul1, culmen length, to cranium (mm). cul2, culmen length, to feathers (mm). width, culmen width (mm). height, culmen height (mm). gape, gape width at the commissures (mm). bmass, body mass (g). gpt, gut passage time (min). Time to first appearance of barium suplphite marker administered orally to the bird. gizz, gizzard mass (g). liver, liver mass (g). intleng, intestine length (from pylorus to cloaca)(mm). trans, transit speed of food throughout the digestive system (mm/min). Estimated as intleng/gpt. source: Sources of data (numerical coding): 1, PJ: Pedro Jordano data. Either measurements of birds in hand or at the Estación Biológica de Doñana (CSIC) collection. 2, Tobias et al. 2022. AVONET: morphological, ecological and geographical data for all birds. Ecology Letters 25:581-597. doi: 10.1111/ele.13898. And 3, Dunning, J. 2008. CRC Handbook of avian body masses. 2nd edition. CRC Press, Boca Raton, USA. (Body mass). Entries with NA indicate no data available for the specific variable.

**Table 3: Suppl. Mat. Table S3.** Avian species included in this study, with indication of fruit feeding while fruit foraging and fruit handling for averaged observations at different plant species. ttot, total time of the observation (s). tpar, total time bird was stopped (not moving or feeding) (s). fru, number of fruits “touched”. swa, number of fruits swallowed. drop, number of fruits “dropped”, usually by a failure to handle in the bill, but also after pecking or tearing-off the pulp. fail, number of fruits failed to be handled and/or ingested, yet not dropped, usually by a failure in detaching the fruit from its peduncle. carry, number of fruits carried away from tree (e.g., in the bill). peck, number of fruits pecked, tearing-off pulp and usually dropping the seed. pick, number of fruits picked. away, number of fruits taken away from tree (swallowed or carried in the bill). beneath, number of fruits estimated to be dropped beneath the plant (usually after pecking the pulp, but also due to handling failures. Entries with dashes indicate no data available for the specific variable.

| species_bird | species_plant | n | ttot | tpar | fru | swa | drop | fail | carry | peck | pick | away | beneath |
| --- | --- | --- | --- | --- | --- | --- | --- | --- | --- | --- | --- | --- | --- |
| *Columba palumbus* | *Prunus mahaleb* | 7 | 239.00±35.57 | 97.29±15.97 | 20.29±3.75 | 20.29±3.75 | 0.00 | 0.00 | 0.00 | 0.00 | 0.00 | 20.29±3.75 | 0.00 |
| *Curruca iberiae* | *Prunus mahaleb* | 9 | 53.67±11.97 | 13.33±1.90 | 3.11±0.84 | 1.67±0.33 | 1.44±0.58 | 0.00 | 0.00 | 0.00 | 0.00 | 1.67±0.33 | 1.44±0.58 |
| *Curruca communis* | *Prunus mahaleb* | 36 | 45.92±3.50 | 17.50±1.53 | 2.83±0.22 | 2.31±0.20 | 0.53±0.14 | 0.00 | 0.00 | 0.00 | 0.00 | 2.31±0.20 | 0.53±0.14 |
| *Curruca melanocephala* | *Pistacia lentiscus* | 1 | 113.00 | 21.00 | 5.00 | 5.00 | --- | --- | --- | --- | --- | --- | --- |
| *Curruca melanocephala* | *Prunus mahaleb* | 3 | 31.67±5.36 | 10.33±1.67 | 1.33±0.33 | 1.00±0.00 | 0.33±0.33 | 0.00 | 0.00 | 0.00 | 0.00 | 1.00±0.00 | 0.33±0.33 |
| *Curruca melanocephala* | *Rubus ulmifolius* | 21 | 37.29±3.37 | --- | 30.38±1.99 | 30.38±1.99 | 0.00 | 0.00 | 0.00 | 0.00 | 0.00 | 30.38±1.99 | --- |
| *Cyanistes caeruleus* | *Prunus mahaleb* | 12 | 69.17±14.53 | 5.67±0.62 | 2.58±0.51 | 0.00 | 0.00 | 0.00 | 0.00 | 0.33±0.26 | 2.25±0.58 | 0.00 | 2.58±0.51 |
| *Cyanistes caeruleus* | *Rubus ulmifolius* | 5 | 82.00±4.67 | --- | 16.80±1.24 | 1.00±0.63 | 0.00 | 0.00 | 0.00 | 15.80±1.28 | 0.00 | 16.80±1.24 | --- |
| *Cyanistes caeruleus* | *Taxus baccata* | 3 | 76.33±4.63 | --- | 1.00±0.00 | 1.00±0.00 | --- | --- | --- | --- | --- | --- | --- |
| *Dendrocopos major* | *Prunus mahaleb* | 4 | 73.50±14.20 | 13.75±7.96 | 4.50±0.96 | 4.25±0.85 | 0.00 | 0.25±0.25 | 0.00 | 0.00 | 0.00 | 4.25±0.85 | 0.00 |
| *Erithacus rubecula* | *Pistacia lentiscus* | 3 | 33.00±4.16 | 16.33±2.19 | 2.00±0.58 | 2.00±0.58 | 0.00 | 0.00 | 0.00 | 0.00 | 0.00 | 2.00±0.58 | 0.00 |
| *Erithacus rubecula* | *Prunus mahaleb* | 13 | 46.92±4.48 | 30.85±3.39 | 2.00±0.28 | 2.00±0.28 | 0.00 | 0.00 | 0.00 | 0.00 | 0.00 | 2.00±0.28 | 0.00 |
| *Erithacus rubecula* | *Rubus ulmifolius* | 11 | 65.18±4.14 | --- | 58.09±1.62 | 58.09±1.62 | 0.00 | 0.00 | 0.00 | 0.00 | 0.00 | 58.09±1.62 | --- |
| *Erithacus rubecula* | *Taxus baccata* | 24 | 66.21±12.19 | --- | 1.04±0.09 | 1.04±0.09 | --- | --- | --- | --- | --- | --- | --- |
| *Ficedula hypoleuca* | *Prunus mahaleb* | 1 | 17.00 | 14.00 | 1.00 | 1.00 | 0.00 | 0.00 | 0.00 | 0.00 | 0.00 | 1.00 | 0.00 |
| *Ficedula hypoleuca* | *Rubus ulmifolius* | 4 | 39.00±1.41 | --- | 19.50±1.26 | 15.50±4.92 | 0.00 | 0.00 | 0.00 | 4.00±4.00 | 0.00 | 19.50±1.26 | --- |
| *Fringilla coelebs* | *Prunus mahaleb* | 51 | 86.84±6.94 | 23.43±2.22 | 3.29±0.22 | 0.00 | 0.00 | 0.00 | 0.02±0.02 | 1.84±0.28 | 1.47±0.19 | 0.02±0.02 | 3.31±0.22 |
| *Garrulus glandarius* | *Prunus mahaleb* | 32 | 106.28±10.90 | 27.88±3.52 | 5.41±0.67 | 0.66±0.40 | 0.03±0.03 | 0.00 | 0.09±0.05 | 0.00 | 4.62±0.66 | 0.75±0.40 | 4.66±0.66 |
| *Lophophanes cristatus* | *Prunus mahaleb* | 1 | 31.00 | 4.00 | 2.00 | 0.00 | 0.00 | 0.00 | 0.00 | 0.00 | 2.00 | 0.00 | 2.00 |
| *Muscicapa striata* | *Prunus mahaleb* | 1 | 154.00 | 113.00 | 1.00 | 1.00 | 0.00 | 0.00 | 0.00 | 0.00 | 0.00 | 1.00 | 0.00 |
| *Parus major* | *Prunus mahaleb* | 5 | 54.60±9.96 | 12.60±3.17 | 2.20±0.37 | 0.00 | 0.00 | 0.00 | 0.00 | 0.00 | 2.20±0.37 | 0.00 | 2.20±0.37 |
| *Periparus ater* | *Prunus mahaleb* | 18 | 47.39±5.93 | 6.17±1.69 | 2.00±0.27 | 0.00 | 0.00 | 0.22±0.17 | 0.22±0.10 | 0.22±0.13 | 1.56±0.26 | 0.22±0.10 | 1.78±0.27 |
| *Periparus ater* | *Taxus baccata* | 2 | 50.50±37.50 | --- | 1.00±1.00 | 1.00±1.00 | --- | --- | --- | --- | --- | --- | --- |
| *Phoenicurus ochruros* | *Prunus mahaleb* | 45 | 45.40±4.92 | 31.42±4.13 | 1.84±0.16 | 1.51±0.11 | 0.22±0.09 | 0.20±0.09 | 0.00 | 0.00 | 0.00 | 1.51±0.11 | 0.22±0.09 |
| *Phoenicurus ochruros* | *Taxus baccata* | 18 | 39.89±5.41 | --- | 1.17±0.09 | 1.17±0.09 | --- | --- | --- | --- | --- | --- | --- |
| *Phoenicurus phoenicurus* | *Prunus mahaleb* | 12 | 52.75±7.43 | 40.08±6.19 | 2.33±0.22 | 2.25±0.22 | 0.08±0.08 | 0.08±0.08 | 0.00 | 0.00 | 0.00 | 2.25±0.22 | 0.08±0.08 |
| *Serinus citrinella* | *Prunus mahaleb* | 2 | 115.50±56.50 | 24.50±14.50 | 6.50±3.50 | 0.00 | 0.00 | 0.00 | 0.00 | 6.50±3.50 | 0.00 | 0.00 | 6.50±3.50 |
| *Serinus serinus* | *Prunus mahaleb* | 1 | 62.00 | 7.00 | 3.00 | 0.00 | 0.00 | 0.00 | 0.00 | 3.00 | 0.00 | 0.00 | 3.00 |
| *Sitta europaea* | *Prunus mahaleb* | 6 | 68.33±18.52 | 12.50±4.84 | 2.50±0.56 | 0.17±0.17 | 0.00 | 0.00 | 0.83±0.31 | 0.00 | 1.67±0.67 | 1.00±0.26 | 1.67±0.67 |
| *Sitta europaea* | *Taxus baccata* | 3 | 65.00±11.00 | --- | 1.00±0.00 | 1.00±0.00 | --- | --- | --- | --- | --- | --- | --- |
| *Sylvia atricapilla* | *Rubus ulmifolius* | 52 | 51.60±2.06 | --- | 49.33±2.91 | 49.33±2.91 | 0.00 | 0.00 | 0.00 | 0.00 | 0.00 | 49.33±2.91 | --- |
| *Sylvia atricapilla* | *Taxus baccata* | 3 | 46.33±36.92 | --- | 1.33±0.33 | 1.33±0.33 | --- | --- | --- | --- | --- | --- | --- |
| *Sylvia borin* | *Prunus mahaleb* | 1 | 30.00 | 9.00 | 4.00 | 4.00 | 0.00 | 0.00 | 0.00 | 0.00 | 0.00 | 4.00 | 0.00 |
| *Sylvia borin* | *Rubus ulmifolius* | 20 | 74.00±3.70 | --- | 90.85±12.87 | 90.85±12.87 | 0.00 | 0.00 | 0.00 | 0.00 | 0.00 | 90.85±12.87 | --- |
| *Turdus iliacus* | *Crataegus monogyna* | 27 | 62.07±4.01 | 25.26±3.01 | 5.30±0.43 | 5.19±0.40 | 0.11±0.08 | 0.00 | 0.00 | 0.00 | 0.00 | 5.19±0.40 | 0.00 |
| *Turdus merula* | *Crataegus monogyna* | 2 | 58.00±35.00 | --- | 7.00±4.00 | 7.00±4.00 | 0.00 | 0.00 | 0.00 | 0.00 | 0.00 | 7.00±4.00 | 0.00 |
| *Turdus merula* | *Prunus mahaleb* | 15 | 55.20±8.43 | 19.60±4.38 | 6.40±0.61 | 5.93±0.64 | 0.13±0.09 | 0.27±0.15 | 0.20±0.20 | 0.00 | 0.00 | 6.13±0.61 | 0.13±0.09 |
| *Turdus merula* | *Rubus ulmifolius* | 4 | 138.25±27.05 | --- | 120.00±9.77 | 120.00±9.77 | 0.00 | 0.00 | 0.00 | 0.00 | 0.00 | 120.00±9.77 | --- |
| *Turdus merula* | *Taxus baccata* | 13 | 102.00±22.99 | --- | 3.46±0.55 | 3.46±0.55 | --- | --- | --- | --- | --- | --- | --- |
| *Turdus philomelos* | *Taxus baccata* | 10 | 197.20±45.08 | --- | 4.70±0.70 | 4.60±0.73 | 1.00 | --- | --- | --- | --- | --- | --- |
| *Turdus pilaris* | *Crataegus monogyna* | 6 | 83.83±28.69 | 41.60±12.50 | 8.67±2.23 | 8.67±2.23 | 0.00 | 0.00 | 0.00 | 0.00 | 0.00 | 8.67±2.23 | 0.00 |
| *Turdus pilaris* | *Rosa canina* | 1 | 115.00 | 48.00 | 13.00 | 13.00 | 0.00 | 0.00 | 0.00 | 0.00 | 0.00 | 13.00 | 0.00 |
| *Turdus torquatus* | *Berberis vulgaris* | 4 | 81.00±12.90 | 23.75±6.61 | 13.00±2.08 | 13.00±2.08 | 0.00 | 0.00 | 0.00 | 0.00 | 0.00 | 13.00±2.08 | 0.00 |
| *Turdus torquatus* | *Crataegus monogyna* | 55 | 87.44±5.57 | 30.85±2.95 | 9.09±0.59 | 8.60±0.57 | 0.42±0.10 | 0.07±0.04 | 0.00 | 0.00 | 0.00 | 8.60±0.57 | 0.35±0.09 |
| *Turdus torquatus* | *Juniperus communis* | 3 | 63.67±6.39 | 26.67±9.13 | 7.00±0.58 | 7.00±0.58 | 0.00 | 0.00 | 0.00 | 0.00 | 0.00 | 7.00±0.58 | 0.00 |
| *Turdus torquatus* | *Juniperus phoenicea* | 56 | 86.12±7.16 | 25.44±1.97 | 9.88±0.86 | 7.38±0.65 | 1.70±0.21 | 1.02±0.20 | 0.00 | 0.00 | 0.00 | 7.38±0.65 | 2.05±0.27 |
| *Turdus torquatus* | *Lonicera arborea* | 6 | 42.33±8.80 | 11.50±4.78 | 8.00±1.84 | 6.67±1.78 | 0.33±0.21 | 1.50±0.81 | 0.00 | 0.00 | 0.00 | 6.17±1.94 | 0.17±0.17 |
| *Turdus torquatus* | *Rosa canina* | 3 | 92.67±22.92 | 27.00±13.58 | 7.67±0.67 | 7.33±0.33 | 0.33±0.33 | 0.00 | 0.00 | 0.00 | 0.00 | 7.33±0.33 | 0.33±0.33 |
| *Turdus viscivorus* | *Crataegus monogyna* | 2 | 72.50±6.50 | 43.00 | 6.50±1.50 | 6.50±1.50 | 0.00 | 0.00 | 0.00 | 0.00 | 0.00 | 6.50±1.50 | 0.00 |
| *Turdus viscivorus* | *Lonicera arborea* | 2 | 89.00±0.00 | 51.00±0.00 | 12.00±0.00 | 9.00±0.00 | 0.50±0.50 | 2.50±0.50 | 0.00 | 0.00 | 0.00 | 9.00±0.00 | 2.00±1.00 |
| *Turdus viscivorus* | *Prunus mahaleb* | 56 | 80.05±5.00 | 27.84±2.82 | 9.04±0.53 | 9.02±0.52 | 0.02±0.02 | 0.00 | 0.00 | 0.00 | 0.00 | 9.02±0.52 | 0.02±0.02 |
| *Turdus viscivorus* | *Taxus baccata* | 13 | 86.46±22.81 | 29.43±13.29 | 4.77±1.08 | 4.00±0.67 | 0.00 | 1.43±0.87 | 0.00 | 0.00 | 0.00 | 8.57±4.25 | 0.00 |

**Table 4: Suppl. Mat. Table S4.** Avian species included in this study, with indication of per-fruit and per-visit feeding rates for averaged observations at different plant species. ttot.fru, total time/fruit. tpar.fru, total stoped time/fruit. mov.fru, number of moves/fruit taken. fru.min, no. fruits handled/min. seeds.visit, no. seeds estimated to be taken away from tree (dispersed) per visit. Entries with dashes indicate no data available for the specific variable.

| species_bird | species_plant | n | ttot.fru | tpar.fru | mov.fru | fru.min | seeds.vis |
| --- | --- | --- | --- | --- | --- | --- | --- |
| *Columba palumbus* | *Prunus mahaleb* | 7 | 13.90±2.61 | 7.04±2.23 | 0.26±0.03 | 5.36±1.03 | 22.40±4.55 |
| *Curruca iberiae* | *Prunus mahaleb* | 9 | 21.53±2.75 | 7.18±1.94 | 3.44±0.64 | 3.41±0.69 | 1.62±0.37 |
| *Curruca communis* | *Prunus mahaleb* | 36 | 17.83±1.18 | 7.35±0.78 | 2.64±0.16 | 3.83±0.23 | 2.25±0.20 |
| *Curruca melanocephala* | *Pistacia lentiscus* | 1 | 22.60 | 0.19 | 2.40 | 2.70 | 5.00 |
| *Curruca melanocephala* | *Prunus mahaleb* | 3 | 24.67±2.33 | 8.33±1.86 | 7.33±2.40 | 2.50±0.23 | 1.00 |
| *Curruca melanocephala* | *Rubus ulmifolius* | 21 | 1.20±0.07 | --- | --- | 53.16±2.99 | 36.14±1.77 |
| *Cyanistes caeruleus* | *Prunus mahaleb* | 12 | 29.80±4.33 | 2.72±0.36 | 5.68±1.38 | 2.52±0.39 | 0.00 |
| *Cyanistes caeruleus* | *Rubus ulmifolius* | 5 | 4.95±0.35 | --- | --- | 12.36±0.92 | 1.00±0.63 |
| *Cyanistes caeruleus* | *Taxus baccata* | 3 | 76.33±4.63 | --- | --- | 0.80±0.06 | 1.00 |
| *Dendrocopos major* | *Prunus mahaleb* | 4 | 16.94±1.39 | 3.00±1.24 | 2.19±0.49 | 3.62±0.34 | 4.50±0.50 |
| *Erithacus rubecula* | *Pistacia lentiscus* | 3 | 20.17±7.42 | 0.52±0.12 | 3.06±0.34 | 3.70±1.00 | 2.00±0.58 |
| *Erithacus rubecula* | *Prunus mahaleb* | 13 | 26.44±2.35 | 17.47±2.00 | 2.60±0.35 | 2.53±0.26 | 1.90±0.28 |
| *Erithacus rubecula* | *Rubus ulmifolius* | 11 | 1.14±0.09 | --- | --- | 56.62±5.32 | 57.71±2.35 |
| *Erithacus rubecula* | *Taxus baccata* | 22 | 57.55±10.55 | --- | --- | 2.20±0.52 | 1.00±0.10 |
| *Ficedula hypoleuca* | *Prunus mahaleb* | 1 | 17.00 | 14.00 | 2.00 | 3.50 | 1.00 |
| *Ficedula hypoleuca* | *Rubus ulmifolius* | 4 | 0.06±0.02 | --- | --- | 1.27±1.02 | 19.50±1.26 |
| *Fringilla coelebs* | *Prunus mahaleb* | 51 | 28.61±2.17 | 7.90±0.77 | 1.81±0.15 | 2.65±0.17 | 0.03±0.03 |
| *Garrulus glandarius* | *Prunus mahaleb* | 32 | 24.00±2.26 | 7.24±0.93 | 2.04±0.18 | 3.07±0.21 | 1.13±0.72 |
| *Lophophanes cristatus* | *Prunus mahaleb* | 1 | 15.50 | 2.00 | 1.50 | 3.90 | 1.00 |
| *Muscicapa striata* | *Prunus mahaleb* | 1 | 154.00 | 113.00 | 2.00 | 0.40 | 0.00 |
| *Parus major* | *Prunus mahaleb* | 5 | 26.93±5.33 | 5.87±1.08 | 4.00±1.05 | 2.54±0.42 | 0.00 |
| *Periparus ater* | *Prunus mahaleb* | 18 | 27.96±4.66 | 2.84±0.40 | 3.02±0.26 | 2.87±0.37 | 0.25±0.11 |
| *Periparus ater* | *Taxus baccata* | 1 | 44.00 | --- | --- | 0.70±0.70 | 0.00 |
| *Phoenicurus ochruros* | *Prunus mahaleb* | 45 | 28.35±3.25 | 20.64±3.03 | 1.69±0.09 | 3.12±0.29 | 1.48±0.11 |
| *Phoenicurus ochruros* | *Taxus baccata* | 18 | 37.03±5.74 | --- | --- | 2.56±0.51 | 1.21±0.11 |
| *Phoenicurus phoenicurus* | *Prunus mahaleb* | 12 | 23.37±3.41 | 17.84±2.90 | 1.85±0.15 | 3.16±0.40 | 2.30±0.26 |
| *Serinus citrinella* | *Prunus mahaleb* | 2 | 18.44±1.24 | 3.62±0.28 | 1.06±0.26 | 3.30±0.20 | 0.00 |
| *Serinus serinus* | *Prunus mahaleb* | 1 | 20.67 | 2.33 | 1.00 | 2.90 | 0.00 |
| *Sitta europaea* | *Prunus mahaleb* | 6 | 28.12±4.73 | 5.25±1.77 | 4.33±1.57 | 2.48±0.42 | 1.00±0.26 |
| *Sitta europaea* | *Taxus baccata* | 2 | 65.00±11.00 | --- | --- | 0.95±0.15 | 1.00 |
| *Sylvia atricapilla* | *Rubus ulmifolius* | 52 | 1.12±0.04 | --- | --- | 58.30±2.48 | --- |
| *Sylvia atricapilla* | *Taxus baccata* | 3 | 44.00±38.00 | --- | --- | 7.03±3.41 | 1.00±0.21 |
| *Sylvia borin* | *Prunus mahaleb* | 1 | 7.50 | 2.25 | 2.25 | 8.00 | --- |
| *Sylvia borin* | *Rubus ulmifolius* | 20 | 1.26±0.20 | --- | --- | 78.73±12.99 | 92.90±17.71 |
| *Turdus iliacus* | *Crataegus monogyna* | 27 | 13.06±1.07 | 0.39±0.03 | 0.38±0.05 | 5.71±0.58 | 6.00±0.63 |
| *Turdus merula* | *Crataegus monogyna* | 2 | 8.06±0.39 | --- | 0.82±0.18 | 7.45±0.35 | --- |
| *Turdus merula* | *Prunus mahaleb* | 15 | 8.78±0.99 | 3.13±0.57 | 0.72±0.09 | 8.21±0.95 | 6.09±0.80 |
| *Turdus merula* | *Rubus ulmifolius* | 4 | 1.13±0.13 | --- | --- | 55.23±5.81 | 120.00±9.77 |
| *Turdus merula* | *Taxus baccata* | 12 | 33.63±9.24 | --- | --- | 3.25±0.87 | 3.46±0.55 |
| *Turdus philomelos* | *Taxus baccata* | 10 | 40.03±9.46 | --- | --- | 3.05±1.10 | 4.33±1.05 |
| *Turdus pilaris* | *Crataegus monogyna* | 6 | 8.70±0.83 | 0.46±0.03 | 0.67±0.08 | 7.22±0.66 | 7.80±2.06 |
| *Turdus pilaris* | *Rosa canina* | 1 | 8.85 | 0.42 | 0.31 | 6.80 | 15.00 |
| *Turdus torquatus* | *Berberis vulgaris* | 4 | 6.58±1.28 | 0.28±0.04 | 0.38±0.08 | 10.15±1.87 | 11.75±2.59 |
| *Turdus torquatus* | *Crataegus monogyna* | 55 | 10.36±0.48 | 0.31±0.02 | 0.55±0.04 | 6.58±0.33 | 9.64±0.69 |
| *Turdus torquatus* | *Juniperus communis* | 3 | 9.17±0.98 | 0.40±0.10 | 1.31±0.11 | 6.70±0.76 | 9.00±1.53 |
| *Turdus torquatus* | *Juniperus phoenicea* | 56 | 9.62±0.45 | 0.31±0.07 | 0.59±0.05 | 6.99±0.33 | 10.00±1.03 |
| *Turdus torquatus* | *Lonicera arborea* | 6 | 7.23±2.69 | 0.26±0.11 | 1.16±0.33 | 12.13±2.37 | 24.67±7.76 |
| *Turdus torquatus* | *Rosa canina* | 3 | 12.05±2.98 | 0.28±0.10 | 0.54±0.16 | 5.73±1.63 | 9.33±1.86 |
| *Turdus viscivorus* | *Crataegus monogyna* | 2 | 11.54±1.66 | 0.54 | 0.55±0.05 | 5.30±0.80 | 8.00 |
| *Turdus viscivorus* | *Lonicera arborea* | 2 | 7.42 | 0.57 | 1.12±0.04 | 8.10 | 36.00 |
| *Turdus viscivorus* | *Prunus mahaleb* | 56 | 9.56±0.56 | 3.44±0.40 | 0.59±0.05 | 7.39±0.40 | 9.70±0.58 |
| *Turdus viscivorus* | *Taxus baccata* | 13 | 20.44±5.40 | 0.49±0.12 | 1.71±0.43 | 5.68±1.11 | 6.73±2.74 |

**Table 5: Suppl. Mat. Table S5.** Avian species included in this study, with indication of fruit foraging data (use frequency of different manoeuvres and moves) and fruit handling for averaged observations at different plant species. pct.swa, percentage of fruit feeding attempts ending in swallowed fruit. pct.fails, percentage of fruit feeding attempts ending in failed fruit picking. pct.drop, percentage of fruit feeding attempts ending in dropped fruit. pct.peck, percentage of fruit feeding attempts pecking the fruit for pulp. pct.pick, percentage of fruit feeding attempts ending in picking fruit. pct.away, percentage of fruit feeding attempts ending in fruit moved away from plant. pct.beneath, percentage of fruit feeding attempts ending in fruit being dropped beneath the canopy. branch, number of fruits taken while perched from branch. jump, number of fruits taken while jumping. hang, number of fruits taken while hanging from branch. mov, number of moves while foraging in the tree. front, number of fruits taken to the front of perch. down, number of fruits taken below the perched position. up, number of fruits taken above the perched position. Entries with dashes indicate no data available for the specific variable.

| Species bird | Species plant | n | pct.swa | pct.fails | pct.drop | pct.peck | pct.pick | pct.away | pct.beneath | branch | jump | hang | mov | front | down | up |
| --- | --- | --- | --- | --- | --- | --- | --- | --- | --- | --- | --- | --- | --- | --- | --- | --- |
| *Columba palumbus* | *Prunus mahaleb* | 7 | 100.00±0.00 | 0.00 | 0.00 | 0.00 | 0.00 | 100.00±0.00 | 0.00 | 20.29±3.75 | 0.00 | 0.00 | 5.71±1.60 | 0.00 | --- | --- |
| *Curruca iberiae* | *Prunus mahaleb* | 9 | 72.78±10.96 | 0.00 | 27.22±10.96 | 0.00 | 0.00 | 72.78±10.96 | 27.22±10.96 | 2.67±0.60 | 0.00 | 0.00 | 7.78±1.22 | 2.78±0.64 | --- | 1.00±0.03 |
| *Curruca communis* | *Prunus mahaleb* | 36 | 84.77±3.85 | 0.00 | 15.23±3.85 | 0.00 | 0.00 | 84.77±3.85 | 15.23±3.85 | 2.83±0.22 | 0.00 | 0.00 | 7.14±0.65 | 2.83±0.22 | --- | --- |
| *Curruca melanocephala* | *Pistacia lentiscus* | 1 | 100.00 | 0.00 | 0.00 | 0.00 | 0.00 | 100.00 | 0.00 | 5.00 | 0.00 | 0.00 | 12.00 | 4.00 | --- | --- |
| *Curruca melanocephala* | *Prunus mahaleb* | 3 | 83.33±16.67 | 0.00 | 16.67±16.67 | 0.00 | 0.00 | 83.33±16.67 | 16.67±16.67 | 1.33±0.33 | 0.00 | 0.00 | 11.33±6.36 | 1.33±0.33 | --- | --- |
| *Curruca melanocephala* | *Rubus ulmifolius* | 21 | 100.00±0.00 | 0.00 | 0.00 | 0.00 | 0.00 | 100.00±0.00 | --- | --- | --- | --- | --- | --- | --- | --- |
| *Cyanistes caeruleus* | *Prunus mahaleb* | 12 | 0.00 | 0.00 | 0.00 | 16.67±11.24 | 83.33±11.24 | 0.00 | 100.00±0.00 | 2.58±0.51 | 0.00 | 0.00 | 11.25±2.06 | 2.00±0.37 | --- | --- |
| *Cyanistes caeruleus* | *Rubus ulmifolius* | 5 | 5.90±3.98 | --- | --- | 94.10±3.98 | --- | 5.80±3.95 | --- | --- | --- | --- | --- | --- | --- | --- |
| *Cyanistes caeruleus* | *Taxus baccata* | 3 | 100.00±0.00 | --- | --- | --- | --- | --- | --- | --- | --- | --- | --- | --- | --- | --- |
| *Dendrocopos major* | *Prunus mahaleb* | 4 | 95.83±4.18 | 4.18±4.18 | 0.00 | 0.00 | 0.00 | 95.83±4.18 | 0.00 | 4.50±0.96 | 0.25±0.25 | 1.50±0.50 | 8.50±0.96 | 1.50±1.50 | --- | 0.08±0.08 |
| *Erithacus rubecula* | *Pistacia lentiscus* | 3 | 100.00±0.00 | 0.00 | 0.00 | 0.00 | 0.00 | 100.00±0.00 | 0.00 | 1.00±0.58 | 1.00±0.58 | 0.00 | 6.33±2.40 | 2.00±0.58 | --- | --- |
| *Erithacus rubecula* | *Prunus mahaleb* | 13 | 100.00±0.00 | 0.00 | 0.00 | 0.00 | 0.00 | 100.00±0.00 | 0.00 | 0.46±0.22 | 1.69±0.31 | 0.00 | 4.46±0.43 | 1.92±0.26 | --- | --- |
| *Erithacus rubecula* | *Rubus ulmifolius* | 11 | 100.00±0.00 | 0.00 | 0.00 | 0.00 | 0.00 | 100.00±0.00 | --- | --- | --- | --- | --- | --- | --- | --- |
| *Erithacus rubecula* | *Taxus baccata* | 22 | 100.00±0.00 | --- | --- | --- | --- | --- | --- | 0.00 | 1.00 | 0.00 | --- | 0.00 | --- | --- |
| *Ficedula hypoleuca* | *Prunus mahaleb* | 1 | 100.00 | 0.00 | 0.00 | 0.00 | 0.00 | 100.00 | 0.00 | 0.00 | 1.00 | 0.00 | 2.00 | 1.00 | --- | --- |
| *Ficedula hypoleuca* | *Rubus ulmifolius* | 4 | 76.47±23.52 | --- | --- | 94.10 | --- | 76.25±23.75 | --- | --- | --- | --- | --- | --- | 0.31±0.13 | 0.09±0.05 |
| *Fringilla coelebs* | *Prunus mahaleb* | 51 | 0.00 | 0.00 | 0.00 | 47.45±6.30 | 53.72±6.21 | 0.39±0.39 | 91.18±2.13 | 3.27±0.21 | 0.00 | 0.00 | 5.49±0.52 | 2.00±0.30 | --- | --- |
| *Garrulus glandarius* | *Prunus mahaleb* | 32 | 6.84±4.18 | 0.00 | 0.26±0.26 | 0.00 | 85.87±5.91 | 13.87±5.80 | 86.13±5.80 | 5.38±0.68 | 0.03±0.03 | 0.00 | 9.19±1.21 | 2.47±0.64 | --- | --- |
| *Lophophanes cristatus* | *Prunus mahaleb* | 1 | 0.00 | 0.00 | 0.00 | 0.00 | 100.00 | 0.00 | 100.00 | 2.00 | 0.00 | 0.00 | 3.00 | 2.00 | --- | --- |
| *Muscicapa striata* | *Prunus mahaleb* | 1 | 100.00 | 0.00 | 0.00 | 0.00 | 0.00 | 100.00 | 0.00 | 0.00 | 1.00 | 0.00 | 2.00 | 1.00 | --- | --- |
| *Parus major* | *Prunus mahaleb* | 5 | 0.00 | 0.00 | 0.00 | 0.00 | 100.00±0.00 | 0.00 | 100.00±0.00 | 2.20±0.37 | 0.00 | 0.00 | 8.40±2.20 | 2.20±0.37 | --- | --- |
| *Periparus ater* | *Prunus mahaleb* | 18 | 0.00 | 8.33±6.06 | 0.00 | 9.44±6.08 | 76.21±8.94 | 14.35±7.68 | 85.65±7.68 | 2.00±0.27 | 0.00 | 0.06±0.06 | 5.67±0.76 | 1.39±0.34 | --- | 0.11±0.06 |
| *Periparus ater* | *Taxus baccata* | 2 | 0.00 | 0.00 | 100.0 | 100.00 | 0.00 | 0.00 | 100.00 | 100.0 | 0.00 | 0.00 | 0.00 | 100.0 | 0.00 | 0.00 |
| *Phoenicurus ochruros* | *Prunus mahaleb* | 45 | 88.85±3.68 | 8.89±3.78 | 6.70±2.67 | 0.00 | 0.00 | 88.85±3.68 | 6.70±2.67 | 0.73±0.14 | 1.07±0.13 | 0.00 | 2.89±0.26 | 1.67±0.13 | --- | 0.17±0.11 |
| *Phoenicurus ochruros* | *Taxus baccata* | 18 | 100.00±0.00 | --- | --- | --- | --- | --- | --- | 0.00 | 1.00 | 0.00 | --- | 0.00 | --- | --- |
| *Phoenicurus phoenicurus* | *Prunus mahaleb* | 12 | 97.22±2.78 | 2.78±2.78 | 2.78±2.78 | 0.00 | 0.00 | 97.22±2.78 | 2.78±2.78 | 1.08±0.34 | 1.25±0.33 | 0.00 | 4.25±0.46 | 2.08±0.26 | --- | --- |
| *Serinus citrinella* | *Prunus mahaleb* | 2 | 0.00 | 0.00 | 0.00 | 100.00±0.00 | 0.00 | 0.00 | 100.00±0.00 | 6.50±3.50 | 0.00 | 0.00 | 6.00±2.00 | 6.50±3.50 | --- | --- |
| *Serinus serinus* | *Prunus mahaleb* | 1 | 0.00 | 0.00 | 0.00 | 100.00 | 0.00 | 0.00 | 100.00 | 3.00 | 0.00 | 0.00 | 3.00 | 3.00 | --- | --- |
| *Sitta europaea* | *Prunus mahaleb* | 6 | 4.17±4.17 | 0.00 | 0.00 | 0.00 | 56.95±18.81 | 59.72±18.56 | 56.95±18.81 | 2.50±0.56 | 0.00 | 0.00 | 12.50±7.15 | 2.50±0.56 | --- | --- |
| *Sitta europaea* | *Taxus baccata* | 3 | --- | --- | --- | 100.00±0.00 | --- | --- | --- | --- | --- | --- | --- | --- | --- | --- |
| *Sylvia atricapilla* | *Rubus ulmifolius* | 52 | 100.00±0.00 | 0.00 | 0.00 | 0.00 | 0.00 | 100.00±0.00 | --- | --- | --- | --- | --- | --- | --- | --- |
| *Sylvia atricapilla* | *Taxus baccata* | 3 | 100.00±0.00 | --- | --- | --- | --- | --- | --- | --- | --- | --- | --- | --- | --- | --- |
| *Sylvia borin* | *Prunus mahaleb* | 1 | 100.00 | 0.00 | 0.00 | 0.00 | 0.00 | 100.00 | 0.00 | 4.00 | 0.00 | 0.00 | 9.00 | 4.00 | --- | --- |
| *Sylvia borin* | *Rubus ulmifolius* | 20 | 100.00±0.00 | --- | --- | --- | --- | 100.00±0.00 | --- | --- | --- | --- | --- | --- | 1.89±0.36 | 0.70±0.25 |
| *Turdus iliacus* | *Crataegus monogyna* | 27 | 98.77±0.91 | 0.00 | 1.23±0.91 | 0.00 | 0.00 | 98.77±0.91 | 0.00 | 5.26±0.44 | 0.04±0.04 | 0.00 | 2.22±0.39 | 2.81±0.32 | 2.50±1.50 | --- |
| *Turdus merula* | *Crataegus monogyna* | 2 | 100.00±0.00 | 0.00 | 0.00 | 0.00 | 0.00 | 100.00±0.00 | 0.00 | 7.00±4.00 | 0.00 | 0.00 | 5.00±2.00 | 4.50±2.50 | 1.00±0.37 | 0.40±0.24 |
| *Turdus merula* | *Prunus mahaleb* | 15 | 92.85±3.75 | 3.60±2.13 | 1.91±1.30 | 0.00 | 0.00 | 96.19±2.19 | 1.91±1.30 | 6.33±0.57 | 0.07±0.07 | 0.00 | 4.67±0.73 | 5.00±0.44 | --- | --- |
| *Turdus merula* | *Rubus ulmifolius* | 4 | 100.00±0.00 | --- | --- | --- | --- | 100.00±0.00 | --- | --- | --- | --- | --- | --- | --- | --- |

**Table 6: Suppl. Mat. Table SX.** Pearson correlation matrix of species traits and fruit feeding behaviour variables.

### 2 Supplementary Material Figures

**Figure 1: SM Figure 1.**
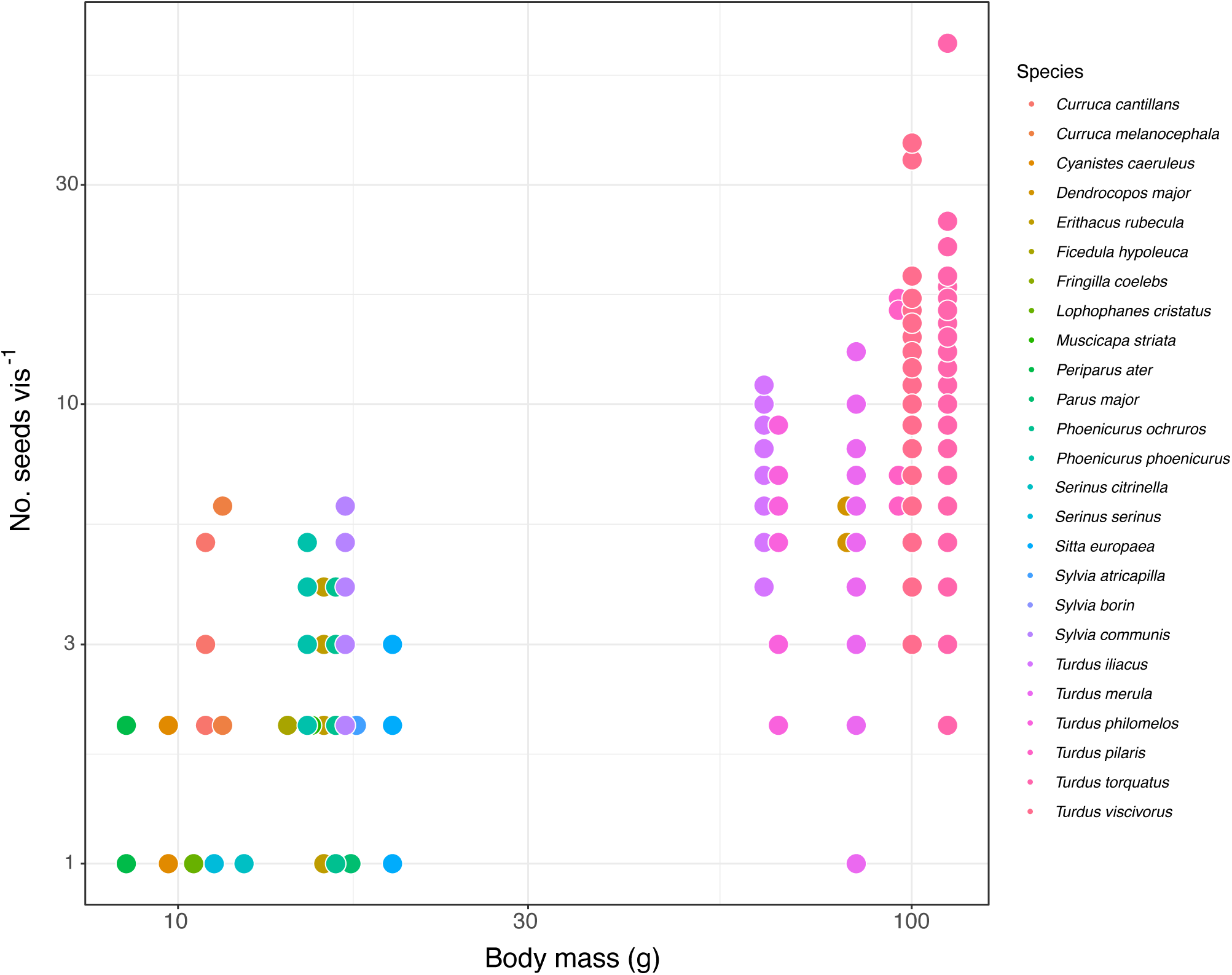
Relationship between body mass (g) and feeding rate per visit (no. seeds estimated to be dispersed successfully from plants in each visit). Points indicate individual foraging observations of different birds species on several plant species (see SM Tables 3, 4.)

**Figure 2: SM Figure 2.**
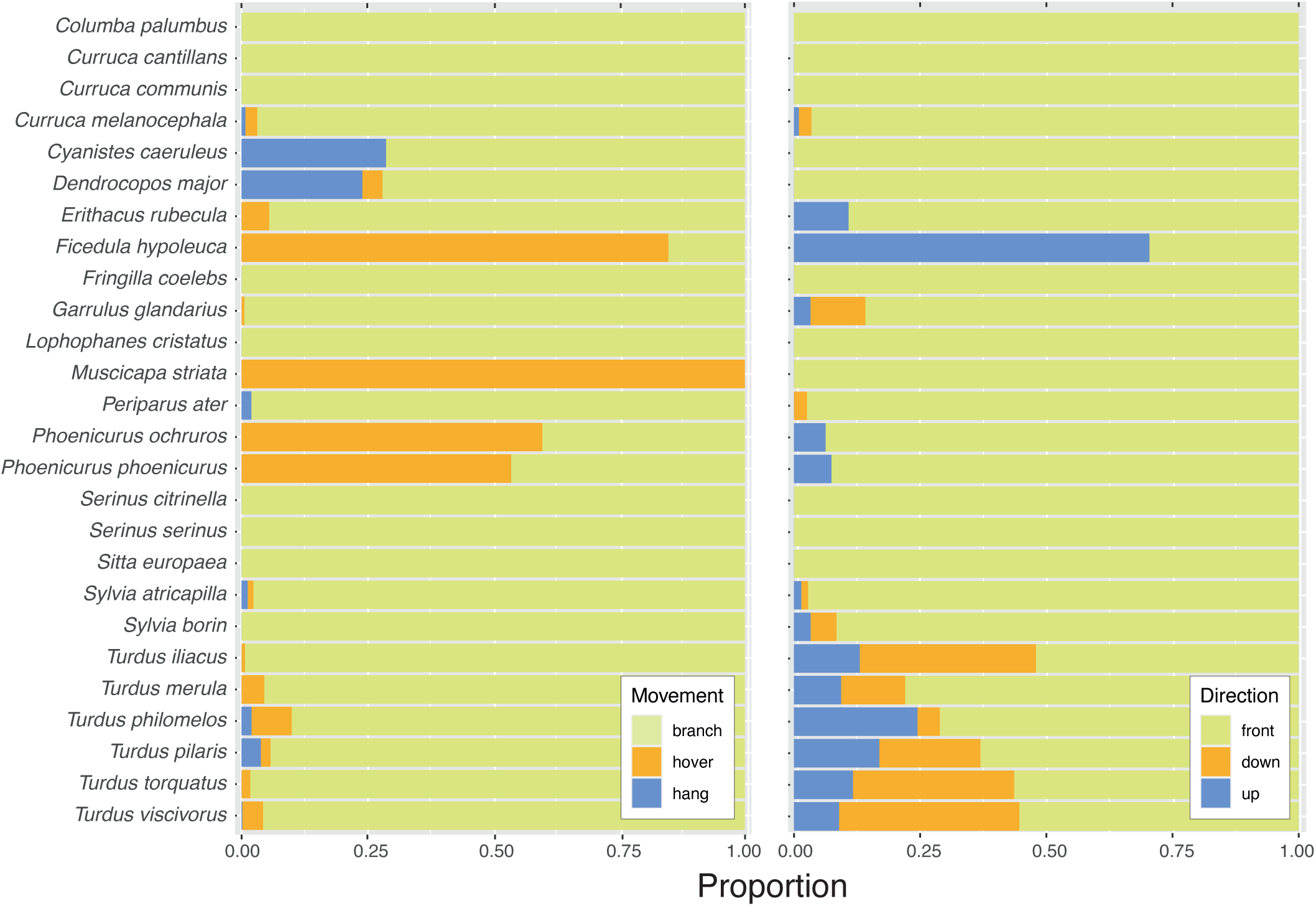
Fruit picking behaviour and fruit accessibility manoeuvres used by foraging avian frugivores.

**Figure 3: SM Figure 3.**
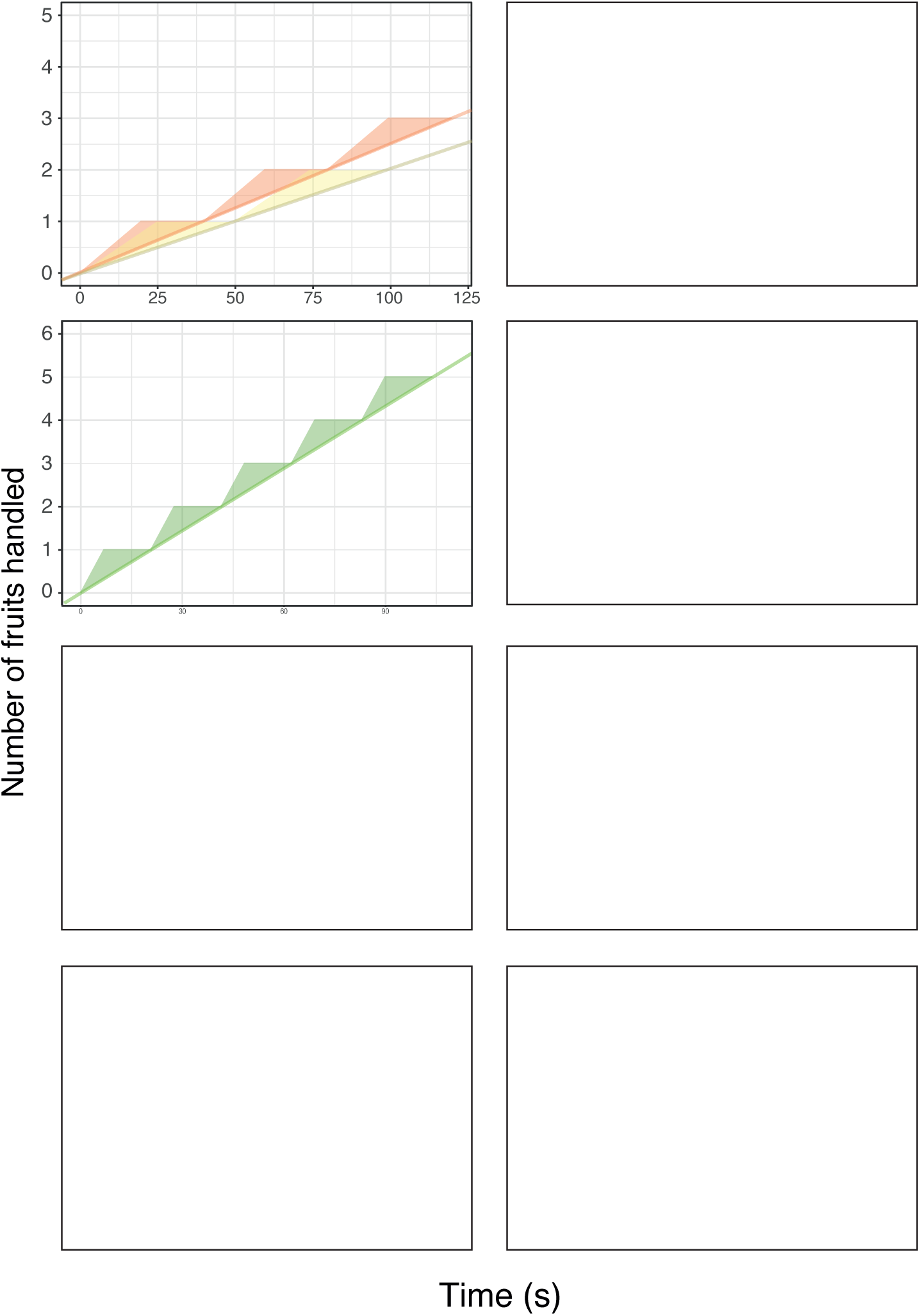
Feeding rates of birds foraging for fruits. The step curves (sawtooth graphs of Cody 1968) show the progression of feeding, with the slope indicating the number of fruits handled per minute. Each step indicates a fruit handled, with the horizontal part of it indicating the time spent stopped and the vertical part of the step encompassing the handling time per fruit. Steep curves are characteristics of efficient foragers, typically SD species (e.g., large *Sylvia*, *Turdus*), and low slopes are cracateristic of PC species (e.g., *P. ater*, *F. coelebs*, *G. glandarius*) that have long handling times while depulping fruits (wide vertical steps). Long horizontal steps are typical of birds taking fruits on the wing while hovering, staying stopped for long periods of time paused by brief movements to pick the fruits while jumping and/or hovering (short vertical steps).

**Figure 4: SM Figure 4.**
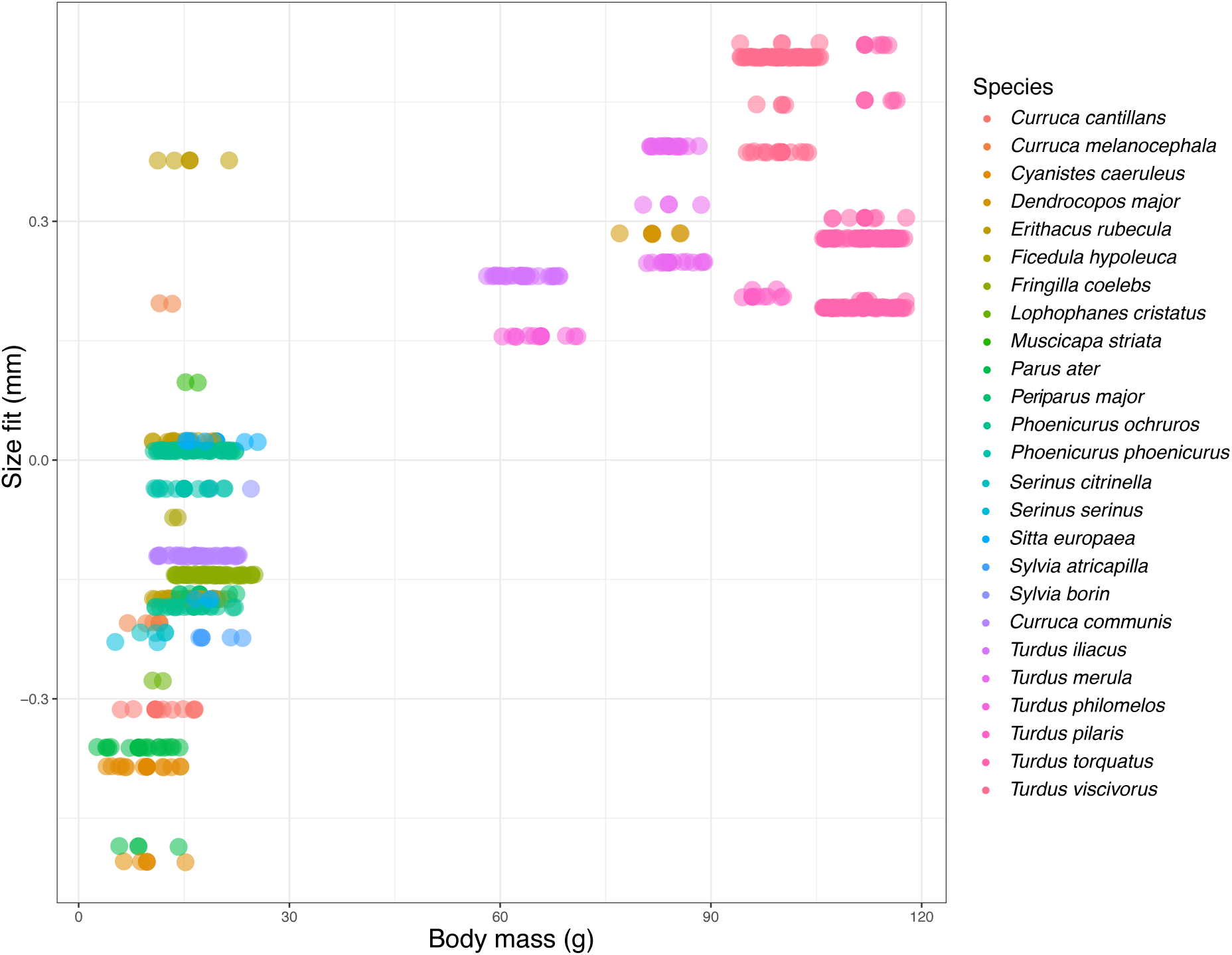
The distribution of size misfit between gape width of avian frugivores and the fruit diameter of their potential food plants in relation with body mass. Each point indicates a pairwise interaction between a frugivore species (color coded) and a plant species. Size fit is computed as the difference between mean fruit size of the plant and mean gape width of the avian frugivore.

